# Mitigation of multi-scale biases in cell-type deconvolution for spatially resolved transcriptomics using HarmoDecon

**DOI:** 10.1101/2024.10.02.616209

**Authors:** Zirui Wang, Ke Xu, Yang Liu, Yu Xu, Lu Zhang

**Affiliations:** Department of Computer Science, Hong Kong Baptist University, Kowloon Tong, Hong Kong

**Keywords:** spatially resolved transcriptomics, deconvolution, sample-level cell-type fractions, dominant cell types, platform effects, multi-scale biases

## Abstract

The advent of spatially resolved transcriptomics (SRT) has revolutionized our understanding of tissue molecular microenvironments by enabling the study of gene expression in its spatial context. However, many SRT platforms lack single-cell resolution, necessitating cell-type deconvolution methods to estimate cell-type proportions in SRT spots. Despite advancements in existing tools, these methods have not addressed biases occurring at three scales: individual spots, entire tissue samples, and discrepancies between SRT and reference scRNA-seq datasets. These biases result in overbalanced cell-type proportions for each spot, mismatched cell-type fractions at the sample level, and data distribution shifts across platforms. To mitigate these biases, we introduce HarmoDecon, a novel semi-supervised deep learning model for spatial cell-type deconvolution. HarmoDecon leverages pseudo-spots derived from scRNA-seq data and employs Gaussian Mixture Graph Convolutional Networks to address the aforementioned issues. Through extensive simulations on multi-cell spots from STARmap and osmFISH, HarmoDecon outperformed 11 state-of-the-art methods. Additionally, when applied to legacy SRT platforms and 10x Visium datasets, HarmoDecon achieved the highest accuracy in spatial domain clustering and maintained strong correlations between cancer marker genes and cancer cells in human breast cancer samples. These results highlight the utility of HarmoDecon in advancing spatial transcriptomics analysis.

## 1 Introduction

Spatially resolved transcriptomics (SRT) is a powerful technology that enables the exploration of gene expression profiles and the spatial landscape of corresponding transcripts. SRT has the potential to uncover the mechanism of cell-cell interactions[1, 2], elucidate tumor microenvironment [3, 4], monitor embryo development [5, 6], and explore the community structure of nerve [7] and immune cells [8]. Nowadays, SRT data can be produced using imaging-based and sequencing-based platforms. Imaging-based SRT platforms are designed using *in situ* hybridization or fluorescence microscopy, such as STARmap [9], seqFISH [10], osmFISH [11], and MERFISH [12]. They could provide gene expression with single-cell resolution and low dropout rates, but their performance is limited by the small numbers of cells and genes that can be captured.

Sequencing-based SRT platforms, including legacy ST [13], Slide-seq [14], 10X Visium [15] and Stereo-seq[16], utilize next-generation sequencing to capture whole transcriptomes across a large number of cells. However, these platforms come with a trade-off: each captured cell-like spatial unit (with a radius of 10-100 µm), known as a spot, may contain multiple cells. This fact leads to low spatial resolution at the cellular level, which substantially impacts the performance of downstream tasks, such as inferring cell-cell interactions[1, 2] and spatial domain clustering[17, 18]. To increase the resolution of sequencing-based SRT techniques (referred to as SRT hereafter), cell-type deconvolution has been introduced to predict the cell-type proportion for each spot using single-cell RNA-seq (scRNA-seq) data from the same tissue. Numerous computational tools have been developed for cell-type deconvolution, which can be categorized into three types: matrix-based approaches, statistics-based approaches, and deep learning-based approaches. Matrix-based approaches utilize non-negative matrix factorization and non-negative least squares for cell-type deconvolution, such as SPOTlight [19], SpatialDWLS [20], CARD [21], and Redeconve [22]. Statistics-based approaches, including Stereoscope [23], RCTD [24], and Cell2location [25], typically assume the transcript read counts follow Poisson distribution or negative binomial distribution. They utilize linear models to predict the parameters of these two distributions by weighted aggregating different cell type signatures. There is a growing trend in developing deep learning-based approaches that generate pseudo-spots using cells from scRNA-seq or SRT platforms with single-cell resolution, such as DSTG [26], Tangram [27], STdGCN [28], and SPACEL [29]. Recent spatially informed cell-type deconvolution approaches, such as CARD and STdGCN, integrate spatial information to ensure that spots in close proximity have similar cell-type proportions.

While these cell-type deconvolution tools have been extensively used across various SRT platforms [30, 2], our preliminary study has identified some unsolved biases. First, spots from most of the platforms are typically small and contain a limited number of cells. For example, in the 10x Genomics Visium platform, each spot has a diameter of 55 *µ*m and generally captures between 1 and 10 cells [15]. Many existing cell-type deconvolution methods for SRT are inherited from the ones designed for bulk RNA-seq data [31], which measures gene expression across hundreds of thousands of cells and may cover all the candidate cell types. These approaches ignore the fact that the cells from different candidate cell types are not evenly distributed for each spot of SRT. We downloaded two SRT datasets with single-cell resolution (STARmap [9] and osmFISH [11]) and aggregated individual cells into multi-cell spots based on their spatial coordinates (see **Methods**, Figure 2a), with each simulated spot typically containing 1 to 10 cells (**Supplementary Fig. 1**). Our observations revealed that cells from different cell types were not uniformly distributed within these simulated multi-cell spots. Instead, there commonly exist dominant cell types, with some types absent in the spot, demonstrating a long-tailed distribution while visualizing their proportions ordered by abundance(Figure 2a and b). In contrast, many existing methods, such as Cell2Location and SPACEL, tended to produce overbalanced cell-type proportions for each spot (**Supplementary Fig. 2 and Fig. 5**).

**Fig. 1:**
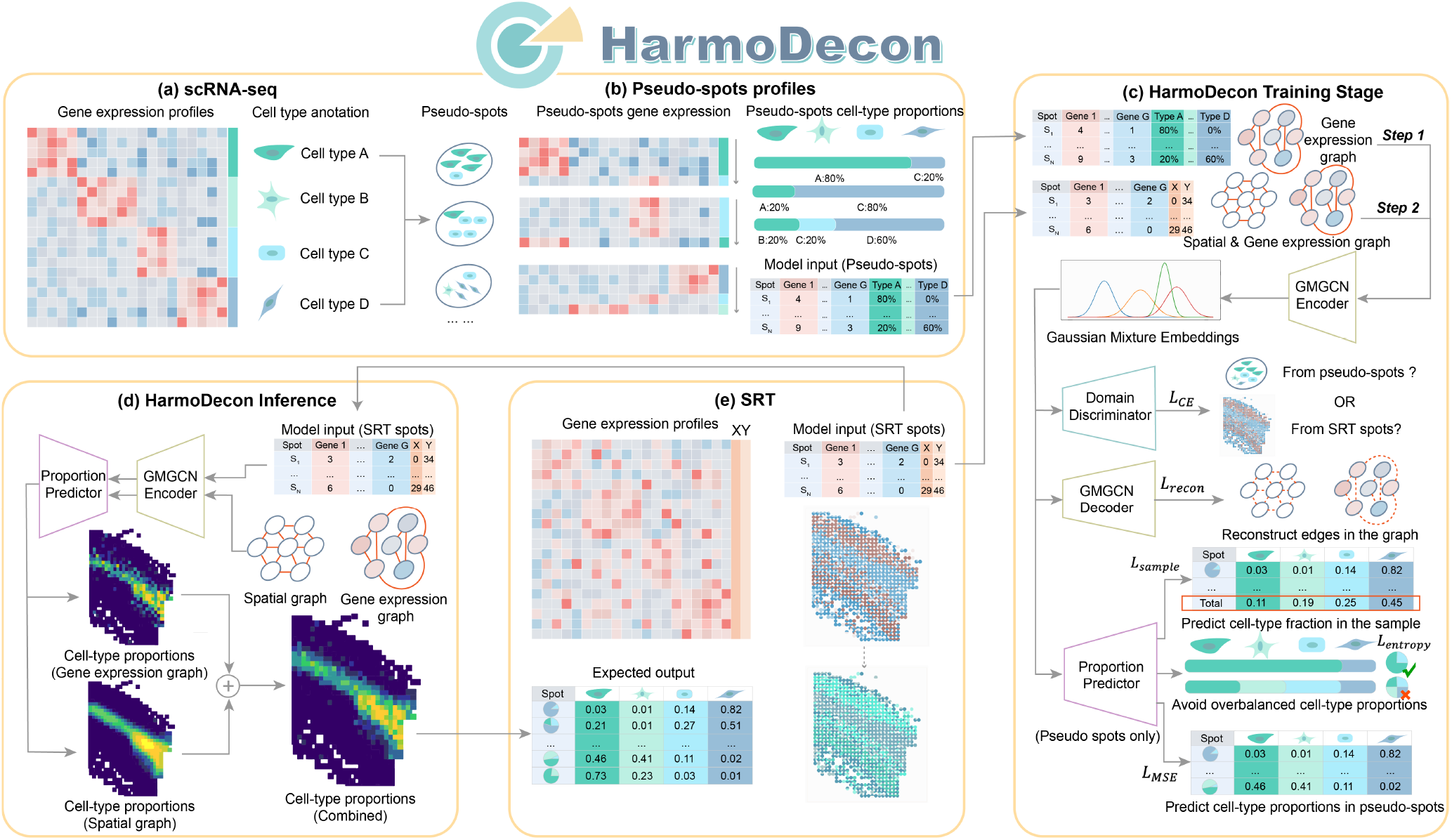
The workflow of HarmoDecon. HarmoDecon is designed for spatially informed cell-type deconvolution for SRT data. HarmoDecon requires 1. gene expression and cell-type annotations of single cells from scRNA-seq (**a**); 2. gene expression and spatial coordinates of SRT spots (**e**); In the training stage, HarmoDecon generates pseudo-spots with known cell-type proportions by randomly selecting cells from scRNA-seq data (**b**). HarmoDecon incorporates GMGCN to accept spatial and gene expression graphs from SRT spots and pseudo-spots (**c**). GMGCN optimizes its parameter by considering five loss functions: 1. a cross-entropy loss function to overcome platform effect (*L*_*CE*_); 2. a reconstruction loss function to reconstruct graph structures (*L*_*recon*_); 3. a sample loss function to minimize the difference between predicted and expected sample-level cell-type fractions (*L*_*sample*_); 4. an entropy-based loss function to avid overbalanced cell-type proportion (*L*_*entropy*_); 5. a MSE loss function to minimize the predicted and expected cell-type proportions for each spot (*L*_*MSE*_). In the inference procedure, HarmoDecon averages cell-type proportions predicted from the spatial and gene expression graphs. Finally, HarmoDecon gives a set of cell-type heat maps and estimates the cell-type proportion of SRT spots.

**Fig. 2:**
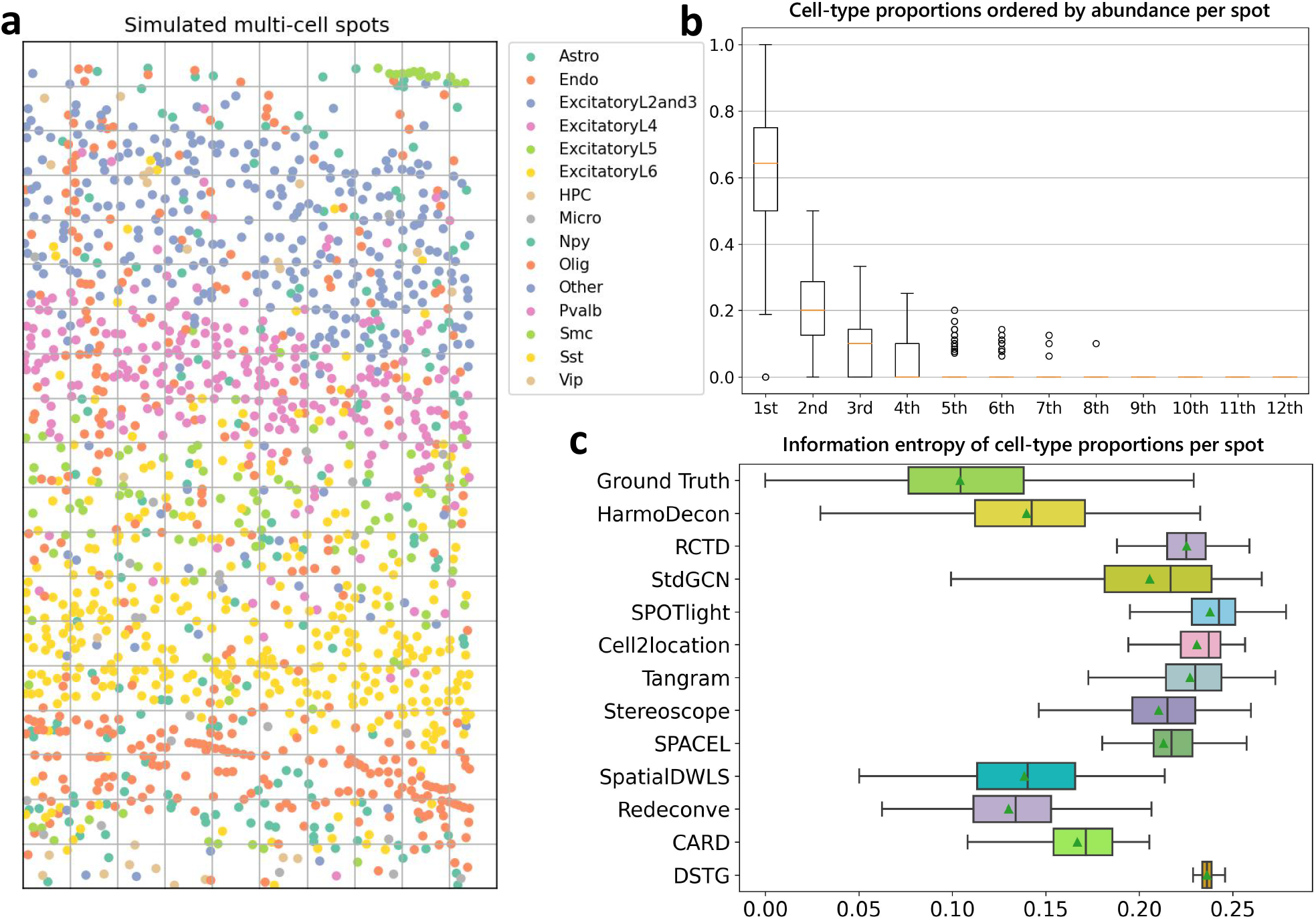
Preliminary studies on the simulated multi-cell spots from the STARmap dataset. **a**. Simulate multi-cell spots on the STARmap dataset. We simulated multi-cell spots by dividing the landscape into quadrilateral grids, each representing a rectangular grid containing 1 to 16 cells. The gene expression profile for each simulated spot was calculated by aggregating the gene expression of all involved cells. Dots with different colors represent single cells from different cell types. **b**. Boxplots of cell-type proportions ordered by abundance per spot (n = 189) in the simulated STARmap data. We sort the number of abundance of each cell type by spot. The i-th box of the plot represents the cell type with the i-th abundance instead of a specific cell type. Three cell types (Other, Npy and HPC) not occurring in the reference scRNA-seq are excluded. **c**. The box plot of information entropy of cell-type proportions predicted by different tools (n = 189). Three cell types (Other, Npy and HPC) are excluded. A lower entropy value indicates a distribution with fewer dominant cell types, while a higher value suggests more balanced cell-type proportions.

Second, the bias from sample-wise cell-type fractions is often overlooked. As we know, similar to spots, each sample slice has its own cell-type fractions that reflect its characteristics. Existing approaches primarily focus on optimizing cell-type proportions at the spot level without ensuring that the predicted sample-wide cell-type fractions are consistent with expected values. In our analysis of simulated multi-cell spot datasets, we calculated the sample-level cell-type fractions by aggregating cell-type proportions across all spots and observed that the sample-level cell-type fractions estimated by existing tools, deviated significantly from the ground truth, especially for dominant cell types (Figure 4 b; **Supplementary Fig. 3**).

Third, a bias can also arise from the platform effect between SRT and scRNA-seq datasets. The platform effect has been documented in previous studies [24] and was also observed in our simulated multi-cell spots (**Supplementary Fig. 4**). This platform effect leads to shifts in gene expression values, complicating downstream analyses. While some statistics-based approaches, such as RCTD, have developed strategies to address this issue, deep learning-based methods have yet to tackle this challenge effectively.

In this study, we propose HarmoDecon (Figure 1), a novel semi-supervised deep learning model designed to mitigate these multi-scale biases in SRT data deconvolution. We generated pseudo-spots with known cell-type proportions by aggregating single cells from scRNA-seq data with annotated cell types (**Methods**, Figure 1 a, b). HarmoDecon incorporates Gaussian Mixture Graph Convolutional Network (GMGCN) to capture the relationship between spots from three graphs (**Methods**): 1. a gene expression similarity graph for pseudo-spots; 2. a gene expression similarity graph for SRTs pots; and 3. a spatial proximity graph for SRT spots. To reduce existing biases, HarmoDecon utilizes 1. an entropy-based loss to avoid overbalanced cell-type distributions across spots, 2. a sample-level loss function to ensure that the predicted cell-type fractions align with the expected proportions from the whole tissue, and 3. a domain adaptation module [32] to mitigate platform effect between SRT and scRNA-seq data (**Methods**). Our extensive benchmarking on various datasets demonstrates that HarmoDecon outperforms 11 state-of-the-art deconvolution tools, offering significant improvements in both spot-level and sample-level predictions.

## 2 Results

### 2.1 The architecture of HarmoDecon and cell-type deconvolution workflow

HarmoDecon is a semi-supervised deep learning model that utilizes GMGCN architecture to perform accurate cell-type deconvolution in sequencing-based SRT data (Figure 1). GMGCN[33] leverages the graph structure to update node features by message passing and assumes the node embeddings follow a Gaussian mixture model. The rationale behind integrating GMGCN into HarmoDecon lies in its inherent ability to capture the spatial and gene expression similarities among SRT spots/pseudo-spots and reflect the fact that SRT spots are from different spatial domains. The model is trained using pseudo-spots generated from scRNA-seq data with known cell-type fractions, and the parameters of GMGCN are optimized through a set of specialized loss functions (**Methods**): 1. Cross-entropy loss: HarmoDecon includes a domain discriminator to differentiate if a given graph derives from scRNA-seq (pseudo-spots) or SRT (SRT spots). By reversing the gradient during backpropagation, the encoder of GMGCN can learn common features for these two platforms to fool the discriminator. This adversarial training design can mitigate the platform effect; 2. Reconstruction loss: HarmoDecon is encouraged to reconstruct the graph structure from node embeddings; 3. Sample loss: HarmoDecon strives to minimize the difference between the predicted and expected sample-level cell-type fractions, which are calculated by averaging the cell-type proportions of all pseudo-spots. 4. Entropy loss: HarmoDecon encourages the prediction of predominant cell types (low information entropy) for SRT spots. 5. HarmoDecon seeks to minimize the mean squared error (MSE) between the predicted and expected cell-type proportions for pseudo-spots. The MSE loss is directly tied to the cell-type deconvolution task and serves as the primary loss function. During inference, HarmoDecon combines predictions from both spatial and gene expression graphs to deliver highly accurate cell-type deconvolution results.

### 2.2 HarmoDecon achieves better cell-type deconvolution on simulated multi-cell spots from STARmap and osmFISH

To simulate SRT spots with known cell-type proportions, we generated synthetic multi-cell spots by aggregating single cells from STARmap [9] and osmFISH [11] technologies (**Methods**, Figure 2). HarmoDecon was benchmarked against 11 existing deconvolution tools—RCTD, STdGCN, SPOTlight, Cell2Location, Tangram, Stereoscope, SPACEL, SpatialDWLS, Redeconve, CARD, and DSTG—using five evaluation metrics: the Pearson correlation coefficient (PCC), structural similarity index measure (SSIM), Jensen-Shannon divergence (JSD), root-mean-square error (RMSE), and a comprehensive rank-based accuracy score (AS). The AS provides a holistic evaluation by considering the ranking of each tool across PCC, SSIM, JSD, and RMSE metrics[34] (**Methods**). The scRNA-seq datasets used by these deconvolution tools were downloaded from the same tissue as those used to generate STARmap [35] (from the mouse visual cortex) and osmFISH [36] (from the mouse somatosensory cortex) datasets. For STARmap, we only kept 12 common cell types that both scRNA-seq and SRT have during evaluation. Three cell types (Other, Npy and HPC) are excluded.

In the simulated multi-cell spots from both STARmap and osmFISH technologies, HarmoDecon outperformed all other state-of-the-art tools across all five evaluation metrics (Figure 3 b and d). HarmoDecon achieved the highest AS values (STARmap: 0.76, osmFISH: 0.77), significantly surpassing the second-best tools: RCTD for STARmap (*p*-value=7.85E-3, **Supplementary Fig. 5**) and Redeconve for osmFISH (*p*-value = 4.88E-6, **Supplementary Fig. 6**). Spatial scatter pie charts showed that the cell-type proportions estimated by HarmoDecon most closely matched the ground truth in the STARmap (Figure 3 a) and osmFISH (Figure 3 c) simulated datasets. In contrast, other methods produced an overbalanced mixture of cell types per spot, which was quite different from the ground truth. The result of HarmoDecon on the STARmap data demonstrated that areas with a high prevalence of eL2/3, eL4, eL5, and eL6 cell types corresponded well with the expected excitatory spatial regions (Figure 3e). In contrast, other methods failed to show clear distributions of these cell types in the spatial domains (**Supplementary Figs 7**). A similar trend was observed with the osmFISH dataset, where HarmoDecon accurately identified regions predominantly composed of L2/3 IT CTX1, L4/5 IT CTX, and L6 CT CTX cell types within their respective L2/3, L4/5, and L6 domains. Other methods struggled to correctly align these cell types with their specific domains (**Supplementary Figs 8**).

**Fig. 3:**
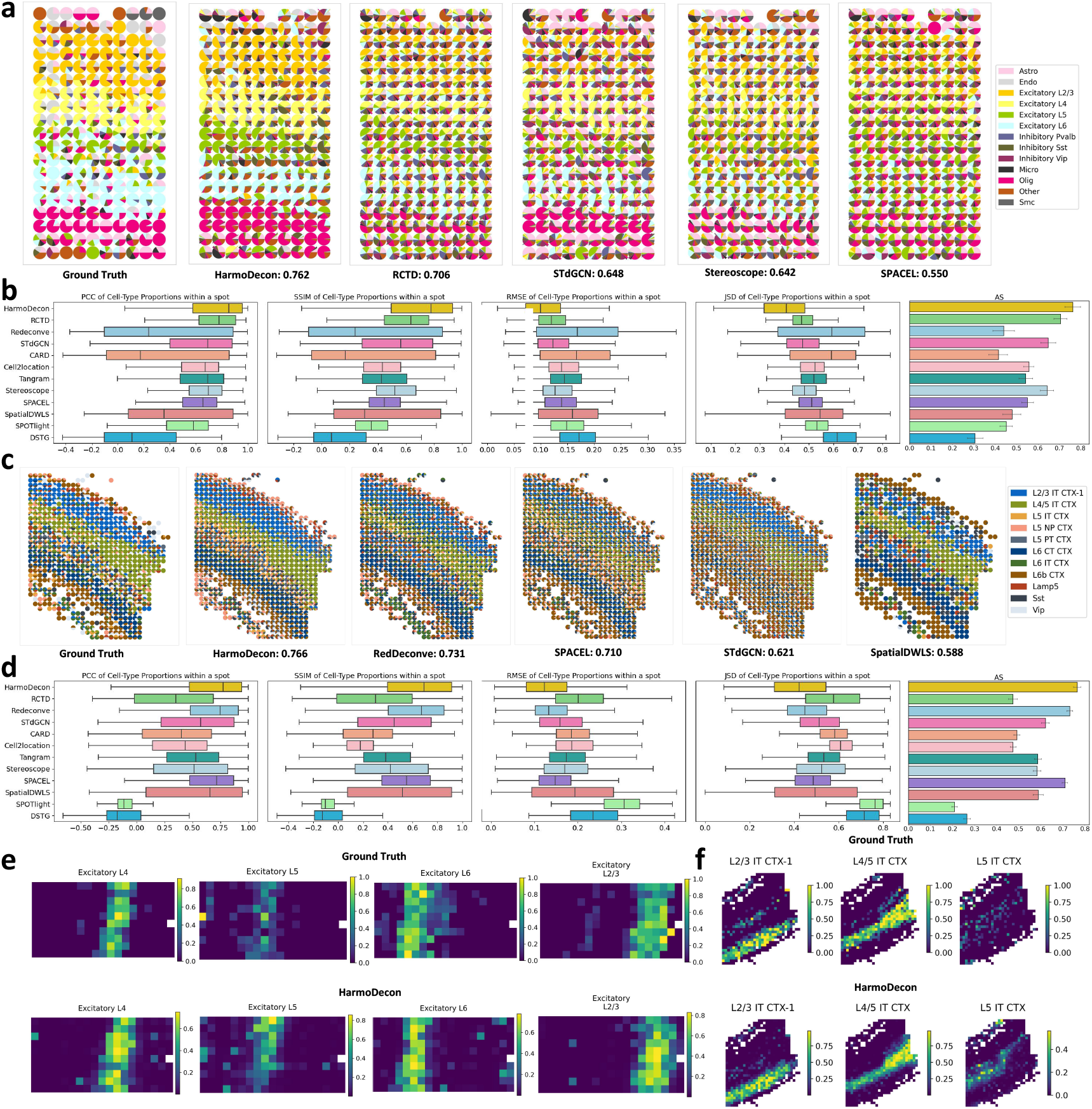
Cell-type deconvolution results on simulated multi-cell spots from STARmap and osmFISH. Spatial scatter pie plots show the cell type proportions for each spot as predicted by different methods on the simulated STARmap (**a**) and osmFISH (**c**) datasets. The corresponding AS is shown below each plot. Boxplots (n = 189 (STARmap), 737 (osmFISH)) of the four evaluation metrics (PCC, SSIM, RMSE, JSD) and bar plots of AS for 12 benchmarking methods on the STARmap (**b**) and osmFISH (**d**) datasets. The bar plots include error bars representing a 95% confidence interval. The proportions of corresponding excitatory neurons predicted by HarmoDecon in the simulated spots from STARmap (**e**) and osmFISH (**f**).

**Fig. 4:**
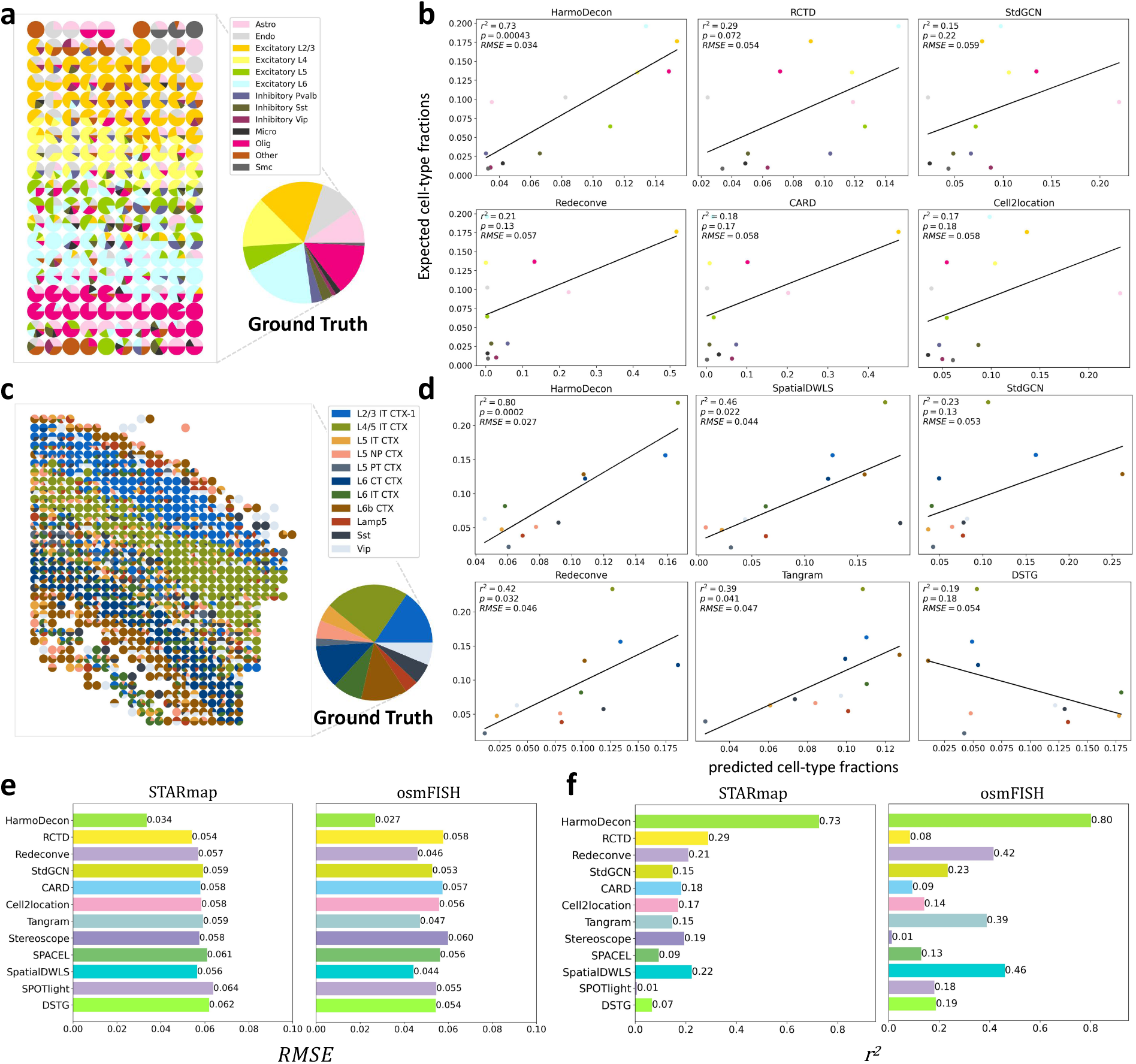
Benchmarking cell-type deconvolution tools for estimating sample-level cell-type fractions. Spatial scatter pie charts illustrate cell-type proportions and the sample-level cell-type fractions (Ground Truth) in simulated datasets from STARmap (**a**) and osmFISH (**c**). Linear regression plots compare the performance of six deconvolution tools with the highest *r*^2^ values in predicting sample-level cell-type fractions for STARmap (**b**) and osmFISH (**d**). The y-axis shows the expected (Ground Truth) fractions, and the x-axis represents the predicted fractions, obtained by averaging proportions across spots. Bar charts compare the performance of different deconvolution tools with respect to RMSE (**e**) and *r*^2^ (**f**).

### 2.3 HarmoDecon captures accurate sample-level cell-type fractions

We calculated the sample-level cell-type fractions for single-cell SRT data from STARmap and osm-FISH (**Methods**, Ground Truth shown in Figure 4 a and c). We observed the sample-level common and rare cell types distributed across different anatomical layers (Figure 4 a and c, left). The sample-level common cell-types dominated the spots in the relevant anatomical layers (STARmap: Excitatory L2/3, Excitatory L4, Excitatory L5, Excitatory L6; osmFISH: L2/3 IT CTX-1, L4/5 IT CTX, L6 CT CTX), while peripheral regions were enriched with rare cell types.

We then performed a linear regression analysis to evaluate the concordance between predicted and ground truth sample-level cell-type fractions of different tools. HarmoDecon demonstrated the highest accuracy, achieving the best *r*^2^ and RMSE values for both STARmap (*r*^2^ =0.73, RMSE=0.034, *p*-value=4E-4 Figure 4 b) and osmFISH (*r*^2^ =0.80, RMSE=0.027, *p*-value=0.0269 Figure 4 d) datasets (**Supplementary Figure 3**). These results substantially outperformed the second-best tools (STARmap: RCTD, *r*^2^=0.29, RMSE=0.054; osmFISH: SpatialDWLS, *r*^2^=0.46, RMSE=0.043). Some large discrepancies between predicted and true cell-type fractions were noted from the results of the other tools. In the STARmap dataset, the most common cell types were Excitatory L2/3, L6, Olig, and L4. Tools like STdGCN and Cell2location notably underestimated the abundances of Excitatory L2/3 and L6, while Redeconve and CARD underestimated Olig and L4. Conversely, RCTD overestimated certain rare cell types, such as Excitatory L5 and Inhibitory Pvalb. A similar pattern emerged with the osmFISH dataset, where STdGCN and Redeconve underestimated the most prevalent cell type, L4/5 IT CTX, while DSTG inverted the overall trend by predicting higher proportions for rare cell types and lower for common ones, resulting in a negative correlation with the true fractions.

### 2.4 HarmoDecon generates predominant cell types for each spot

We employed information entropy to evaluate if the predicted cell-type proportions from deconvolution tools were overbalanced (Figure 2c). A lower entropy value indicates a distribution with fewer dominant cell types, while a higher value suggests more balanced cell-type proportions. Among the methods evaluated, HarmoDecon, SpatialDWLS, and Redeconve yielded entropy values that most closely matched those of the ground truth. Although SpatialDWLS and Redeconve produced lower information entropy, their overall performance was worse than that of HarmoDecon (Figure 3b and **Supplementary Fig. 5**). HarmoDecon effectively inferred dominant cell types for each SRT spot, consistent with our observations in single-cell resolution SRT datasets (**Supplementary Fig. 2**).

### 2.5 HarmoDecon enables accurate spatial domain clustering on legacy SRT datasets

We further evaluated the performance of spatial domain clustering on the cell-type deconvolution results using two SRT datasets from legacy SRT technique: Mouse Olfactory Bulb (MOB) [13] and Human Melanoma [37] datasets. The matched scRNA-seq data were downloaded from two other studies [38, 39]. Spatial domain labels for the spots were assigned by applying K-means clustering to the predicted cell-type proportions, with the value of K determined based on the ground truth. The Mouse Olfactory Bulb (MOB) dataset consists of four anatomical layers: the granule cell layer (GCL), mitral cell layer (MCL), glomerular layer (GL), and olfactory nerve layer (ONL). Compared to other methods (Figure 5 a, **Supplementary Fig. 9**), HarmoDecon accurately reconstructed the original spatial domain structure, achieving a strong alignment with the ground truth (ARI: 0.63, purity: 0.84, Figure 5 c). Competing tools like STdGCN, Redeconve, and RCTD failed to distinguish between the three outer layers (MCL, GL, ONL). While SPACEL, SpatialDWLS, and CARD demonstrated improved clustering performance, they were unable to clearly delineate the boundaries between layers, and their deconvolution results within the same domain showed lower accuracy.

**Fig. 5:**
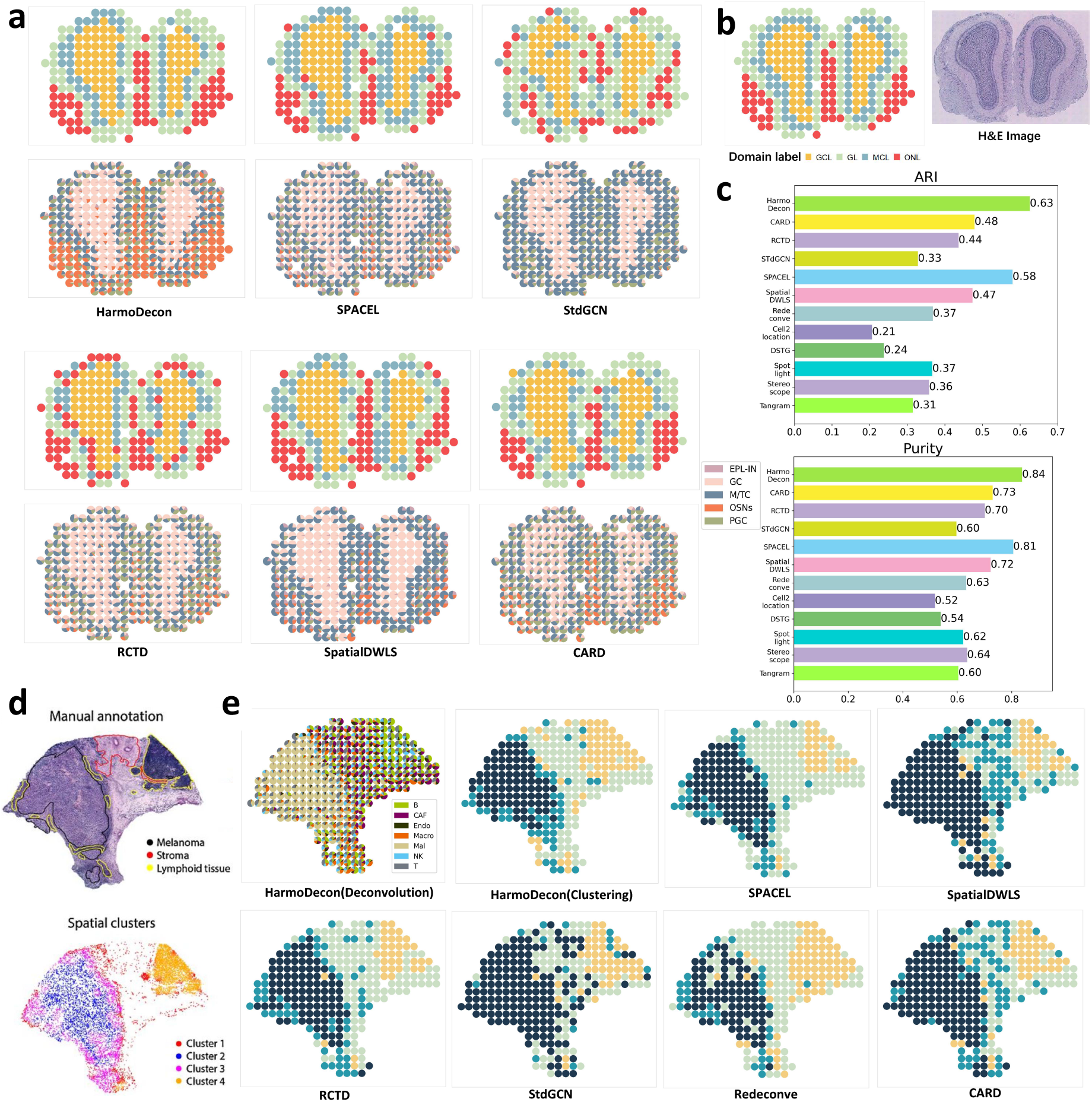
The performance of deconvolution tools on spatial domain clustering. **a**. Spatial domain clustering dot plots and the scatter pie charts of spot cell-type proportions on the MOB dataset. **b**. The ground truth of spatial domain [17] and H&E pathological image [13] of the MOB dataset. **c**. The values of ARI and Purity of spatial domain clustering on the MOB dataset. **d**. The H&E pathological image and spatial domain annotation of the Human Melanoma dataset (credit by [37]). **e**. Spatial domain clustering dot plots show the spatial domains inferred from the results of different deconvolution tools on the Human Melanoma dataset. The scatter pie chart (upper left) shows the cell-type proportions inferred by HarmoDecon.

We found that only HarmoDecon was able to accurately capture the dominant cell types within their corresponding spatial domains. For example, olfactory sensory neurons (OSNs), which are expected to be enriched in the ONL, were predicted by HarmoDecon to make up over 50% of the cell-type proportions across all SRT spots in the ONL (Figure 5 a, in bright orange). Moreover, HarmoDecon was the only method that delineated a distinct boundary for the GL (Figure 5 a, in bright green), with an enrichment of periglomerular cells (PGCs) and mitral/tufted cells (M/TCs). In comparison, the second-best tool, SPACEL, struggled to reconstruct a continuous GL, while the remaining tools frequently misclassified spots as part of the GL.

Based on the H&E image, the Human Melanoma dataset demonstrated four spatial domains (Figure 5 d, spatial clusters): stroma (cluster 1, red), core melanoma (cluster 2, blue), the border area between the lymphoid and tumor tissues(cluster 3, purple) and lymphoid tissue (cluster 4, orange). HarmoDecon, RCTD, and SPACEL outperformed the other tools and could identify the boundary between core melanoma and the border area. RCTD misclassified spots in benign regions, which were primarily composed of cancer cells, leading to an erroneous expansion of the melanoma regions (Figure 5 e, **Supplementary Fig. 10** teal spots). SPACEL curated smaller lymphoid regions, with several spots belonging to the lymphoid tissue (Figure 5 d, manual annotation) assigned to the stroma.

### 2.6 HarmoDecon captures correlations inside cancer regions on 10X Visium SRT datasets

To evaluate the performance of HarmoDecon across different platforms and tissue types, we apply it to two human breast cancer samples sequenced using the 10X Visium technology[15] (BRCA_1: Figure 6c and BRCA_2: Figure 6f). We downloaded scRNA-seq data for matching breast cancer subtypes (BRCA_1: ER+HER2+; BRCA_2: HER2+) and annotated cell types using human breast cancer atlas[40].

**Fig. 6:**
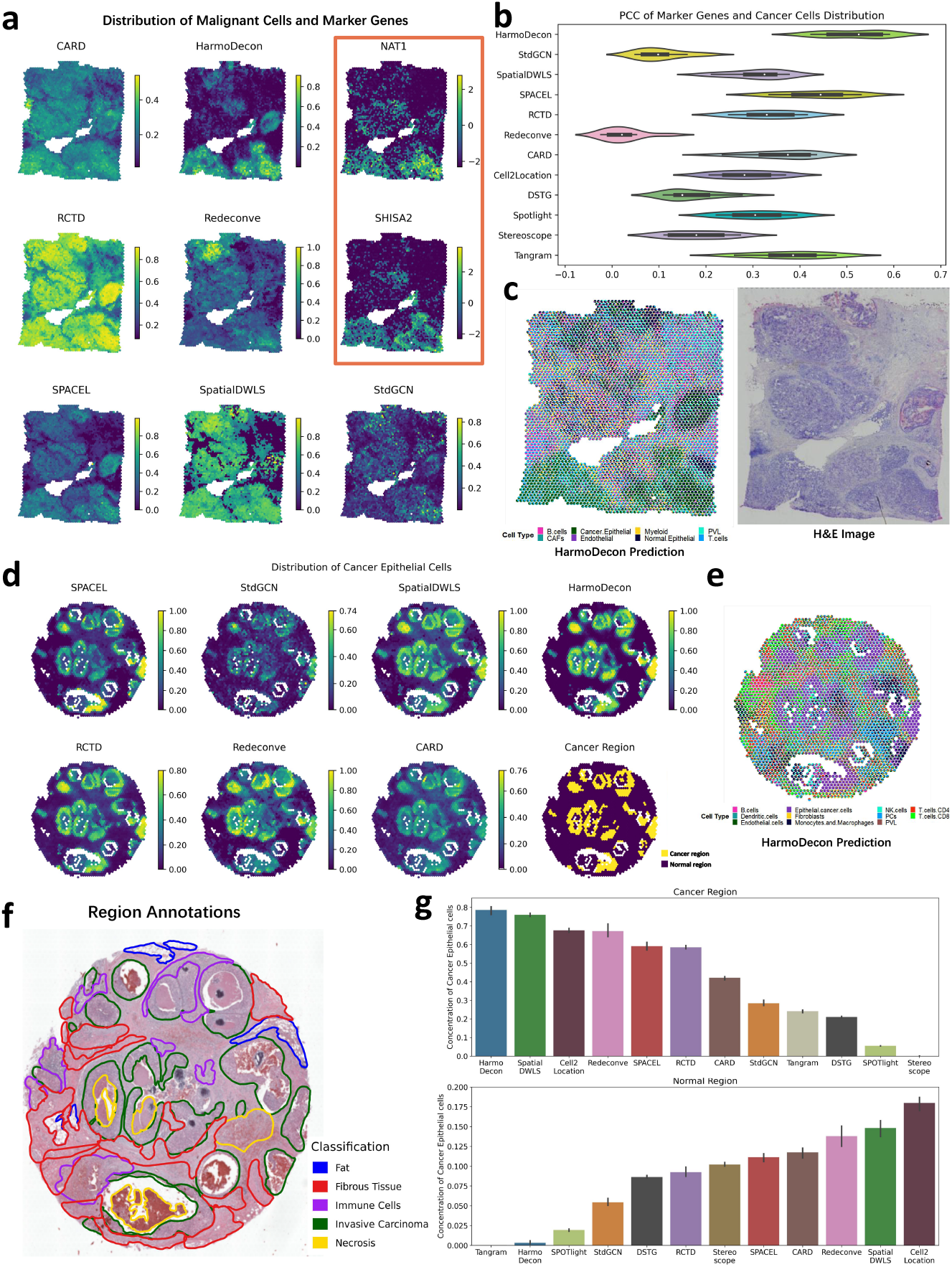
The performance of HarmoDecon on two breast cancer 10X Visium datasets. **a**. Distributions of cancer epithelial cells and gene expression of NAT1 and SHISA2 on the BRCA_1 dataset. **b**. The Violin plot of PCC between the gene expression profiles of ten marker genes and cancer epithelial cell distribution on the BRCA_1 dataset. **c**. The pie chart plots of cell-type proportions predicated by HarmoDecon and the H&E pathological image of the BRCA_1 dataset. **d**. Ground truth of cancer and normal regions and distributions of cancer epithelial cells from different deconvolution methods on the BRCA_2 dataset. **e**. The pie chart of cell-type proportions predicated by HarmoDecon on the BRCA_2 dataset. **f**. Region annotation of the BRCA_2 dataset. **g**. Bar charts of the proportion of cancer epithelial cells in the cancer region (upper) and normal region (lower) inferred by different deconvolution methods in the BRCA_2 dataset.

For the BRCA_1 sample, we examined the colocalization of cancer cells with known cancer marker genes. Ten well-documented breast cancer marker genes were selected (PIP[41], SHISA2[42], PDZK1IP1[43], PLEKHS1[44], NAT1[45], TFAP2C[46], NPNT[47], SCUBE2[48], ASCL2[49], DNAJC1[50]), whose expression is expected to correlate strongly with the presence of cancer epithelial cells. We observed all ten marker genes showed high expression levels in the lower part of the tissue section, with significantly lower expression in the upper part (Figure 6 a: red box and **Supplementary Fig. 11**), suggesting that cancer epithelial cells are also concentrated at the bottom. Only HarmoDecon (Figure 6 a, **Supplementary Fig. 12**) successfully aligned the gene expression patterns of these marker genes with the distribution of cancer epithelial cells, while other methods failed to do so. We also calculated PCC between the gene expression profiles of marker genes and the predicted cancer epithelial cell proportions. HarmoDecon demonstrated the highest PCC, which was significantly better than the second-best tool (SPACEL, p-value=0.002, Figure 6 b). CARD, RCTD, SPACEL and SpatialDWLS tended to assign high proportions of cancer epithelial cells to all spots. Redeconve failed to map the spatial distribution of normal and cancer cells, showing the opposite trend. STdGCN failed to capture any spatial distribution preferences for cancer epithelial cells.

We then investigated if the distribution of cancer epithelial cells matched with the carcinoma regions in the BRCA_2 dataset. Cancer and normal regions were annotated based on the H&E image (**Methods**, Figure 6 d). HarmoDecon, SpatialDWLS, and Redeconve all enriched spots dominated by cancer epithelial cells in the cancerous regions (**Supplementary Fig. 13**). HarmoDecon produced a clear boundary between cancerous and normal regions, with far fewer spots from normal regions predicted to contain cancer epithelial cells (Figure 6 d, **Supplementary Fig. 14**). It also generated the highest median proportion of cancer epithelial cells (0.786) in cancerous regions and the second-lowest median value (0.003) in normal regions. Although Tangram had the smallest median value (0.0002) in normal regions, its median proportion in cancerous regions (0.272) was significantly lower than that of HarmoDecon (Figure 6 g).

## 3 Methods

### 3.1 Data preprocessing

We applied the same data preprocessing strategy to the datasets from SRT and matched scRNA-seq using Scanpy package[51], including 1. library size normalization(*scanpy*.*pp*.*normalize total*), 2. logarithmic normalization on gene expression profiles (*scanpy*.*pp*.*log1p*), 3. gene-wise z-score standardization on gene expression profiles (*scanpy*.*pp*.*scale*), 4. highly variable genes shared in SRT and scRNA-seq (*scanpy*.*pp*.*highly variable genes*, with default thresholds).

### 3.2 Simulate multi-cell spots from STARmap and osmFISH

The landscape of the datasets from STARmap and osmFISH was split into quadrilateral grids with intervals of 750 and 800 pixels. Each single-cell resolution spot (with cell-type annotation) was assigned to its closest grid by computing its distance from grid centers [34, 52]. Each grid represented a simulated multi-cell spot with 1-16 cells (STARmap) and 1-19 cells(osmFISH). A total of 189 (STARmap) and 737 (osmFISH) multi-cell spots are simulated.

### 3.3 Generate pseudo-spots from scRNA-seq data

We used scRNA-seq data to generate pseudo-spots. For each pseudo-spot, we assumed that the number of involved cells (*n*_*c*_) follows a Poisson distribution (with at least one cell per spot), and the number of involved cell types (*n*_*k*_) follows a discrete uniform distribution.

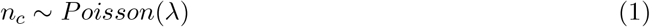

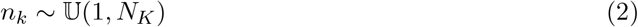

*λ* (*λ* = 5) and *N*_*K*_ (*N*_*K*_ = 4) represent the mean of Poisson distribution and the maximum number of cell types allowed included in a pseudo-spot. For pseudo-spot *i*, we randomly selected *n*_*c*_ cells from *n*_*k*_ cell types with equal probabilities and aggregated their gene raw counts to generate a gene expression profile. The cell-type proportion for the pseudo-spot *i* is represented as *y*_*i*_. In this study, we sampled 50,000 pseudo-spots for each experiment.

### 3.4 Construct spatial and gene expression graphs

During training, HarmoDecon accepts three kinds of graphs: 1. spatial graphs for SRT spots; 2. gene expression graphs for SRT spots; and 3. gene expression graphs for pseudo-spots. The spatial graph connects spots by selecting their top 6 nearest neighbors based on the Euclidean distance of spots’ spatial coordinates. The gene expression graphs are constructed by linking spots (pseudo-spots) to their top nearest neighbors based on the cosine similarities of their gene expression profiles.

### 3.5 Training and inference of HarmoDecon

HarmoDecon is based on the Gaussian Mixture Graph Convolutional Network (GMGCN), which learns representations of spots and pseudo-spots from spatial and gene expression graphs, respectively. For pseudo-spots, *A* ∈ ℝ^*n*×*n*^ is the adjacency matrix of the gene expression graph with gene expression profiles *X* = [*x*_1_, *x*_2_, …, *x*_*n*_] ∈ ℝ^*n*×*g*^ as node feature, where *n* is the number of pseudo-spots in the graph. GMGCN could generate *Z* = [*z*_1_, *z*_2_, …, *z*_*n*_] ∈ ℝ^*n*×512^ as pseudo-spot embeddings, where *z*_*i*_ is sampled from *k*-th component of Gaussian Mixture Model (with means *µ*_*ik*_ and variances *σ*_*ik*_) in the latent space (**Supplementary Notes**). We have 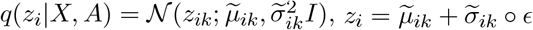, where *ϵ* ∼ 𝒩 (0, *I*)

We define the reconstruction loss function as:

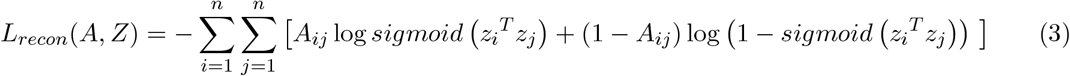

*z*_*i*_ is further passed to a two-layer Multilayer Perceptron (**Supplementary Notes**) to predict the cell-type proportion of pseudo-spot 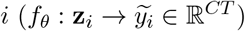. We also introduced mean squared error (MSE) loss to minimize the observed and predicted cell-type proportions.

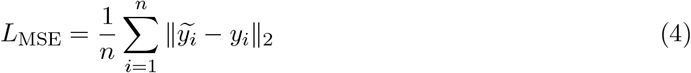

To overcome the platform effect, we utilized a discriminator *f*_*φ*_ : **z**_*i*_ → (**0, 1**) to distinguish if the graph is generated from pseudo-spots or SRT spots. Before the discriminator, there is an additional gradient reversal layer, making the gradient reverse during backpropagation. The final outcome is that the discriminator tries to minimize the domain discriminator loss while the former encoder tries to maximize the domain discriminator loss. In this way, the encoder can extract common features for scRNA-seq and SRT. This process is called adversarial domain adaptation. The true and predicted domain labels of node *i* are represented as *m*_*i*_ and 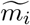 (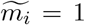 and *m*_*i*_ =1 if the nodes represent SRT spots), respectively. The cross-entropy loss of the domain discriminator is represented as:

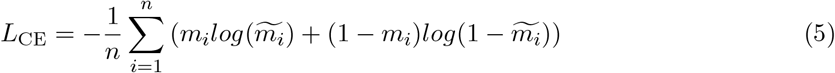

Another task is to learn the whole cell-type fractions at the sample level. We calculated the predicted cell-type fraction by averaging the predicted cell-type proportions of all pseudo-spots in the graph. Similarly, we obtained the true cell-type fraction by approximately averaging the true cell-type proportions of all pseudo-spots (instead of summing by all single cells, see differences in the next section). HarmoDecon encourages minimizing the difference between predicted and true cell-type fractions:

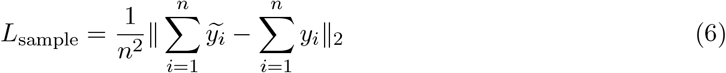

We also employed an entropy-based loss, which penalizes overbalanced cell-type proportions (e.g., 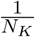) and encourages generating dominant cell type(s) (e.g., *>*50%) for the individual pseudo-spot (first term) and across all pseudo-spots for each batch (second term).

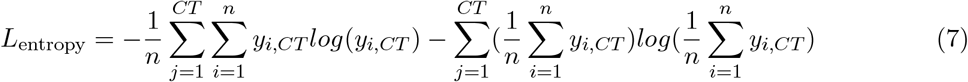

The final loss function on pseudo-spots can be written as *L* = *L*_*recon*_ + 1000 *\* L*_*MSE*_ + *L*_*CE*_ + 1000 *\* L*_*sample*_ + 10 *\* L*_*entropy*_. For the graphs from SRT spots, the final loss function is *L* = *L*_*recon*_ + *L*_*CE*_. In the inference procedure, we provide HarmoDecon with both spatial and gene expression graphs of pseudo-spots to generate 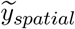 and 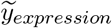. The final cell-type proportion is calculated by

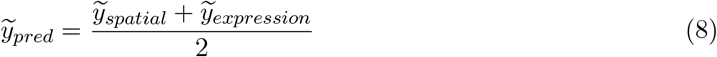

HarmoDecon is trained by Adam optimizer with a learning rate = 0.001 on 20 epochs, each epoch consisting of three types of graphs: 1. 250 gene expression graphs, each with 200 pseudo-spots; 2. a gene expression graph with all SRT spots; 3. a spatial graph with all SRT spots.

### 3.6 Calculation and approximation of sample-level cell-type fractions

In **Results**, we introduce two single-cell resolution SRT datasets to evaluate the performance of deconvolution on the sample level. The “Ground Truth” shown in Figure 4 is calculated by summing all the single cells:

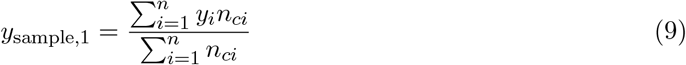

*n* and *n*_*ci*_ represent the number of spots and the number of cells in the spot *i*, respectively. In **Methods**, the sample-level cell-type fraction of the graph built with pseudo-spots is again calculated for *L*_sample_. To simplify it, we calculate the fraction in an approximate way:

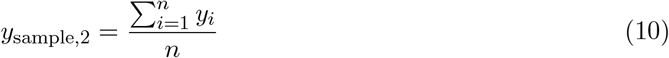

We notice that *y*_sample,1_ = *y*_sample,2_ when *n*_*c*1_ = *n*_*c*2_ = … = *n*_*cn*_. That is, assuming each pseudo-spot has the same number of cells. For STARmap and osmFISH, cell-type fractions obtained by these two approaches have only a small difference in values (**Supplementary Table 1**).

### 3.7 Acquisition of cancer labels of the BRCA dataset

In Figure 6f, we have original manual annotations of the BRCA_2 dataset, which figures out the invasive carcinoma region. To acquire the spot-level annotations of cancer regions, we first exploited a well-curated cell-spot alignment result from [30], in which spots with more than one cancer cell are considered as cancer regions. Then, we adjusted the margin and removed isolated spot points manually to make them more similar to Figure 6f. Finally, we got a cancer region mask like Figure 6d, which contains 532 spots considered as cancer regions and 1986 spots as normal regions. For cancer labels in scRNA-seq dataset, we downloaded scRNA-seq data with matching breast cancer subtypes (BRCA_1: ER+HER2+; BRCA_2: HER2+) and cell types from a human breast cancer atlas[40]. For the ER+HER2+ subtypes in BRCA_1, we referred to GSE176078: CID3586, CID4066, with annotations of normal and cancer epithelial cells in the major cell types provided by the author. While for the BRCA_2, there are only cancer epithelial cells annotated in all three HER2+ subtype samples GSE176078: CID3921, CID45171, CID3838, and we used the minor cell types provided by the author for detailed annotations, following the same criterion as [30].

### 3.8 Compared methods

We compared HarmoDecon to 11 cell-type deconvolution methods: (1) SPOTlight[19] (version 1.4.1), (2) SpatialDWLS[20] (integrated in the R package Giotto, version 1.1.2), (3) Redeconve[22] (version 1.1.0), (4) Stereoscope[23] (integrated in the python package scvi, version 0.6.8), (5) RCTD[24] (integrated in the R package spacexr, version 2.2.1), (6) Cell2location[25] (version 0.1.3), (7) CARD[21] (version 1.14), (8) DSTG[26] (original version in the GitHub page), (9) Tangram[27] (version 1.0.4), (10) STdGCN[28] (original version in the GitHub page), (11) SPACEL[29] (version 1.1.6). For each method, we employed default parameters on their GitHub pages and followed the steps as the tutorials suggested.

### 3.9 Evaluation metrics

For STARmap and osmFISH datasets, we applied five metrics [34] to evaluate the performance of cell-type deconvolution tools. The five metrics are all calculated by spots, so we take a spot *i* as an example:

1. Pearson correlation coefficient(PCC):

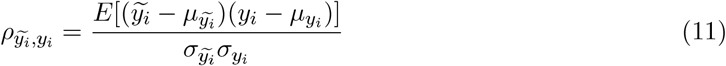

Where

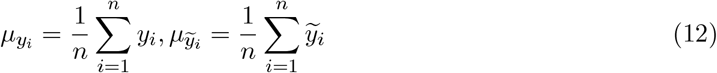

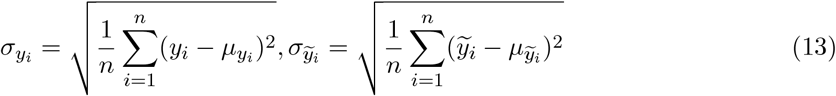

For a given spot, a higher PCC value represents better cell-type deconvolution performance.
2. Structural similarity index measure(SSIM):

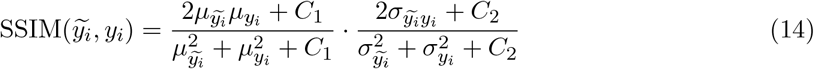

*C*_1_ and *C*_2_ are constants used to improve stability and to avoid division by zero. In this study, we set *C*_1_ and *C*_2_ to 0.01 and 0.03. For a given spot, a higher SSIM value represents better cell-type deconvolution performance.
3. Root-mean-square deviation(RMSE):

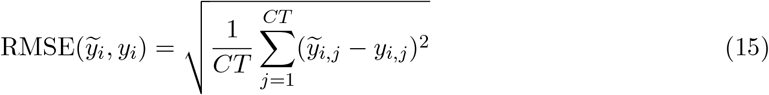

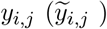 denotes the proportion of cell type *j* in spot *i* from the ground truth (from prediction). For a given spot, a lower RMSE value represents better cell-type deconvolution performance.
4. Jensen–Shannon divergence(JSD):

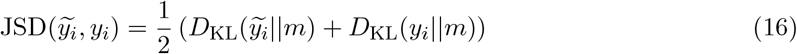

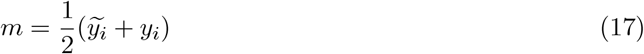

where *D*_KL_ denotes the Kullback-Leibler(KL) divergence. The JS divergence measures the difference between the two distributions, with a lower value indicating greater similarity and better performance on cell-type deconvolution.
5. Accuracy Score(AS):

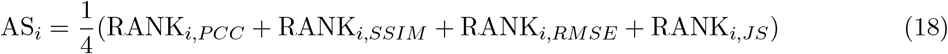

For spot *i*, we sorted the PCC and SSIM in ascending order and sorted RMSE and JS in descending order across the results of 11 cell-type deconvolution methods. RANK_*i,PCC*_, RANK_*i,SSIM*_, RANK_*i,RMSE*_, and RANK_*i,JS*_ refer to the ranks of the candidate method for spot *i* on the PCC, SSIM, RMSE and JS.

For the MOB dataset, we employed Adjusted Rand Index (ARI) and purity to evaluate spatial domain clustering. The ground truth and predicted (by K-Means) spatial domain clustering are denoted as *H* = *{h*_1_, …, *h*_*J*_ *}* and 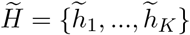, respectively.

We included only the cell types annotated in the scRNA-seq data and excluded any cell types present in the SRT that were not annotated in the scRNA-seq when calculating these metrics.

1. ARI:

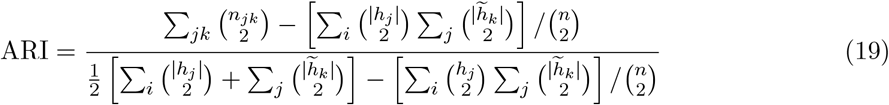

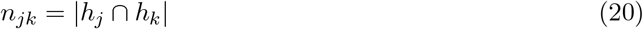
2. Purity:

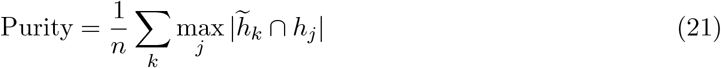

## 4 Discussion

Spatial transcriptomics offers valuable insights into spatial biology, allowing us to understand gene expression profiles in tissue spatial landscapes. Some SRT technologies can not achieve single-cell resolution, and many methods have been developed for cell-type deconvolution. However, despite the advancements of existing tools, our preliminary studies revealed three biases: 1. overbalanced cell-type proportions predicted in individual spots; 2. sample-level cell-type fractions from SRT data fail to align with the expected cell-type fractions; 3. platform effects between scRNA-seq and SRT data; In this study, we introduced HarmoDecon, a novel semi-supervised deep learning model designed for cell-type deconvolution in SRT data. HarmoDecon addresses these three biases by generating pseudo-spots from scRNA-seq data and utilizing GMGCN with specific loss functions. We performed extensive experiments on different SRT technologies and observed that HarmoDecon outperformed 11 state-of-the-art cell-type deconvolution tools.

We performed comprehensive ablation studies to evaluate the significance of the loss functions employed in HarmoDecon (**Supplementary Fig. 15-17**). Among all the loss functions, the entropy loss proved to be the most important factor in improving the performance of HarmoDecon (**Supplementary Fig. 15, w/o_entropy**). In contrast, sample loss (w/o_sample) and domain adaptation loss (w/o_domain) had minor effects in reducing JSD error for cell-type proportions for each SRT spot. The sample loss primarily affects the accuracy of sample-level cell-type fractions rather than spot-level proportions (**Supplementary Fig. 18**). For example, HarmoDecon without the sample loss led to the underestimation of certain cell types (e.g., L6 IT CTX and L6b CTX) in the osmFISH dataset, causing predicted cell-type fractions to deviate from the ground truth. To assess the domain adaptation loss, we performed principal component analysis (PCA) on SRT spots and pseudo-spot latent embeddings from the STARmap and MOB datasets. The results show that domain adaptation loss effectively mitigates batch effects between SRT and scRNA-seq technologies (**Supplementary Fig. 19**).

Additionally, we investigated the influence of hyperparameters on both sampling and training procedures. When building the spatial graph using K-nearest neighbors, we observed that increasing K led to more connections but also introduced errors (**Supplementary Fig. 17, a**). This may be due to oversmoothing, a well-known issue in graph models [53]. Additionally, the model was sensitive to the number of cells (**Supplementary Fig. 16, a**) per pseudo-spots. In the ablation study, we observed that HarmoDecon performed best when the average number of cells across pseudo-spots was close to one across SRT spots.

HarmoDecon employs spatial and gene expression graphs to capture spatial proximity and gene expression similarities between spots. Our results reveal that averaging the cell-type proportions inferred from both graphs leads to significant improvements across all datasets and evaluation metrics tested (**Supplementary Fig. 20**). This strategy is based on the fact that spatially adjacent spots and spots with high gene expression similarities are likely to share similar cell-type proportions. Similar findings have been reported in recent studies, including [28, 21, 30].

To date, only a few deconvolution tools can handle multi-slice SRT data simultaneously. While SPACEL provides a pipeline for 3D SRT data analysis, its integrated deconvolution tool, Spoint, is limited to single-slice data. Currently, HarmoDecon also operates on single-slice SRT data, but we plan to extend its functionality to support multi-slice datasets in the future. We will build spatial and gene expression graphs by considering spatially adjacent spots as well as spots with high gene expression similarities across different slices.

## Supporting information

Supplemental Infomation, including figures and a table.

## 5 Declaration

### 5.1 Ethics approval and consent to participate

Not applicable.

### 5.2 Consent for publication

Not applicable.

### 5.3 Data Availability

The datasets used in this study are all publicly available:

1. Mouse visual cortex (STARmap): https://www.starmapresources.com/data, with matched scRNA-seq from https://portal.brain-map.org/atlases-and-data/rnaseq/mouse-v1-and-alm-smart-seq.
2. Mouse somatosensory cortex (osmFISH): http://linnarssonlab.org/osmFISH, with cell-type labels annotated by [52], matched scRNA-seq from https://portal.brain-map.org/atlases-and-data/rnaseq/mouse-whole-cortex-and-hippocampus-smart-seq SSp Region.
3. Mouse olfactory bulb (Legacy ST): http://www.spatialtranscriptomicsresearch.org, with matched scRNA-seq from GSE121891 in the GEO database.
4. Human melanoma (Legacy ST): https://www.spatialresearch.org/resources-published-datasets/doi-10-1158-0008-5472-can-18-0747/, with matched scRNA-seq from GSE72056 in the GEO database.
5. Human breast cancer ER+/HER2+ (10X Visium): https://support.10xgenomics.com/spatial-gene-expression/datasets/1.1.0/V1_Breast_Cancer_Block_A_Section_1, with matched scRNA-seq from GSE176078: CID3586, CID4066 in the GEO database.
6. Human breast cancer HER2+ (10X Visium): https://cf.10xgenomics.com/samples/spatial-exp/1.3.0/Visium_FFPE_Human_Breast_Cancer/Visium_FFPE_Human_Breast_Cancer_web_summary.html, with matched scRNA-seq from GSE176078: CID3921, CID45171, CID3838 in the GEO database.

### 5.4 Code Availability

The HarmoDecon scripts are available at https://github.com/ericcombiolab/HarmoDecon/tree/main. The custom code for reproducing results in the experiments is available at https://github.com/ericcombiolab/HarmoDecon/tree/main/scripts.

### 5.7 Authors’ contributions

L.Z. conceived and supervised the project. Z.W. and L.Z. designed the method. Z.W. conducted the experiments and implemented HarmoDecon. Z.W. and L.Z. drafted the manuscript. Z.W., L.Z., K.X., and Y.X. discussed the original ideas. Y.L. tested the code and revised the manuscript. All authors reviewed and approved the manuscript.

## 5.8 Acknowledgments

We thank the Research Grant Council (RGC) Hong Kong and Research Committee of Hong Kong for their kind support of this project.

