## Supplemental Infomation, including figures and a table. for "Mitigation of multi-scale biases in cell-type deconvolution for spatially resolved transcriptomics using HarmoDecon"

### Supplemental Notes

#### The structure of HarmoDecon model

HarmoDecon is based on Gaussian Mixture Graph Convolutional Networks (GMGCN). Similar to Gaussian Mixture Variational Autoencoder (GMVAE), our model combines the flexibility of Gaussian mixture models (GMM) with the generative power of variational autoencoders(VAE) to learn complex latent variable representations in high-dimensional data. To enable the model to learn from graph structures, we substitute the dense layers within the encoders with graph convolutional layers, allowing nodes to consider their neighboring nodes in the graph.

Let $A \in R^{n\times n}$ be the adjacency matrix of an unweighted graph (either a spatial graph or a gene expression graph) with node features $X = [x_1, x_2, \dots, x_n] \in R^{n\times g}$ , where $n$ is the number of nodes (spots) and $g$ is the number of features (genes). $Z = [z_1, z_2, \dots, z_n] \in R^{n\times 512}$ is embeddings of $X$ in the hidden layer. $C = [c_1, c_2, \dots, c_n] \in R^{n\times h}$ is a set of one-hot vectors to determine the $i-th$ Gaussian distribution to generate $z_I$ , while $h$ is the number of Gaussian Mixture components.

The latent variable generated by the $i-th$ Gaussian component of GMGCN can be formulated $z_i|c_i \sim N\left(\mu_{c_i}, \sigma_{c_i}^2 I\right)$, with means $\mu_{c_i}$and variances $\sigma_{c_i}^2$. The one-hot vector $c \sim Cate(\pi)$ is sampled from the mixing probability $\pi$, which chooses one component from the Gaussian mixture.

The posterior distribution of the inference model is symbolized as $q(X,A) = N\left(z_i; \tilde{\mu_\iota}, \tilde{\sigma_\iota}^2 I\right)$, $q(X,A) \approx p(z_i) = \frac{\pi_{c_i} p(c_i)}{\sum_{c'_i=1}^{K} \{\pi_{c'_i} p(c'\}_i)}$, $q(X,A) = q(X,A)q(X,A)$.

The parameters of Gaussian distribution $[\tilde{\mu}, log\tilde{\sigma}^2]$ are generated by using a two-layer graph convolutional network (GCN) based encoder as follows:

$$\tilde{\mu} = GCN\left(ReLU\left(GCN\left(X, A; W^{(0)}\right)\right), A; W_{\tilde{\mu}}\right)$$
$$log\tilde{\sigma}^2 = GCN\left(ReLU\left(GCN\left(X, A; W^{(0)}\right)\right), A; W_{\tilde{\sigma}}\right)$$

The latent variable $z_i$ is obtained from $q(X,A) = N\left(z_i; \tilde{\mu_\iota}, \tilde{\sigma}_i{}^2 I\right)$ using the reparameterization trick $z_i = \tilde{\mu_\iota} + \tilde{\sigma_\iota} \circ \epsilon$, where $\epsilon \sim N(0, I)$. For node $i$ and node $j$, we suppose the edge between $i$ and $j$ can be represented by $Z$ as $A_{ij} \mid Z \sim Ber\left(sigmoid\left(z_i^T z_j\right)\right)$, through which we design the decoder and make the model reconstruct the edges.

For the deconvolution task, $z_i$ is further passed to a two-layer Multilayer Perceptron (MLP), projected to the dimension of cell-type number. We define $f_\theta: z_i \rightarrow \tilde{y_\iota} \in R^{CT}$, where $CT$ represents the number of cell types.

$$\tilde{y_\iota} = f_\theta\left(z_i; W_{\theta_1}, b_{\theta_1}, W_{\theta_2}, b_{\theta_2}\right) = softmax\left(W_{\theta_2}\left(W_{\theta_1 z_i} + b_{\theta_1}\right) + b_{\theta_2}\right)$$

The predicted cell-type vector $\tilde{y_\iota}$ is used to calculate mean square loss.

To alleviate the platform effect, we deployed an adversarial domain adaptation module. $z_i$is passed to a one-layer MLP $f_\varphi: z_i \rightarrow (0,1)$ called discriminator. The discriminator tries to make the model distinguish the data sources (from scRNA-seq or SRT) and extract distinct features for the two platforms. However, we want the encoder to extract common features of two platforms and project them into an overlapping latent space. To achieve so, we employed a gradient reversal layer before the discriminator:

$$R(z_i) = z_i$$

$$\frac{dR}{dz_i} = -1$$

During backpropagation, the model presents an inverse gradient for the parameters of the encoder, making it more likely to extract common features to trick the discriminator.

The discriminator is formulated as:

$$m_i = f_\varphi(z_i; W_\varphi, b) = sigmoid(W_{\varphi z_i} + b)$$

The domain labels $m_i$ are then used to calculate the cross-entropy loss.

### Supplemental Table

| **osmFISH** | by cells | by spots |
|---|---|---|
| L2/3 IT CTX-1 | 0.156242 | 0.160234 |
| L4/5 IT CTX | 0.233447 | 0.190375 |
| L5 IT CTX | 0.04737 | 0.052223 |
| L5 NP CTX | 0.051034 | 0.053651 |
| L5 PT CTX | 0.021722 | 0.023797 |
| L6 CT CTX | 0.121958 | 0.09716 |
| L6 IT CTX | 0.081654 | 0.07115 |
| L6b CTX | 0.128239 | 0.157939 |
| Lamp5 | 0.03821 | 0.045849 |
| Sst | 0.057315 | 0.06644 |
| Vip | 0.062811 | 0.081182 |

| **STARmap** | by cells | by spots |
|---|---|---|
| Astro | 0.114336 | 0.119129 |
| Endo | 0.107941 | 0.108231 |
| Excitatory L2/3 | 0.178712 | 0.179833 |
| Excitatory L4 | 0.107702 | 0.104822 |
| Excitatory L5 | 0.076246 | 0.074693 |
| Excitatory L6 | 0.184083 | 0.179567 |
| Inhibitory Pvalb | 0.03124 | 0.03172 |
| Inhibitory Sst | 0.025254 | 0.025475 |
| Inhibitory Vip | 0.012116 | 0.01416 |
| Micro | 0.014047 | 0.01416 |
| Olig | 0.13596 | 0.135964 |
| Smc | 0.012363 | 0.012246 |

**Supplementary Table 1 The sample-level cell-type fraction of osmFISH and STARmap, calculated by counting single cells and by averaging spots. Some ambiguous cell types(e.g. Others) are removed.**

### Supplemental Figures

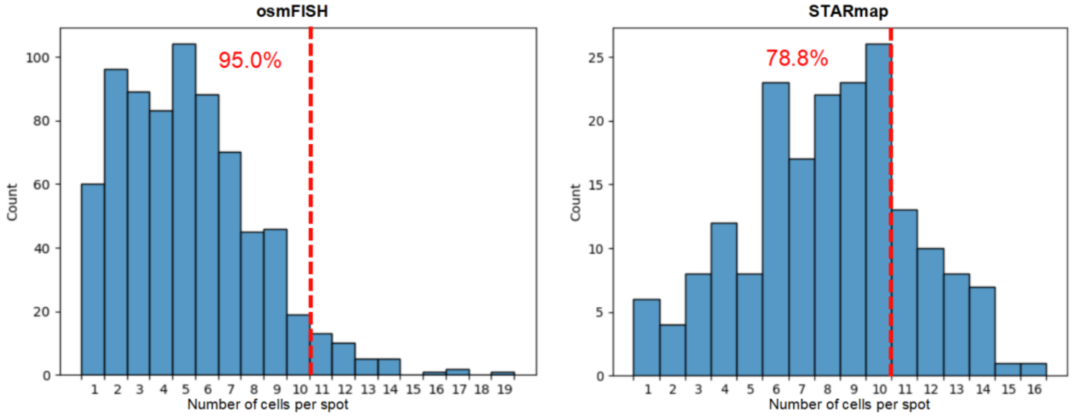


**Supplementary Figure 1. Histogram of number of cells per spot in simulated multi-spots datasets.** Most simulated multi-cell spots contain 1 to 10 cells. STARmap generally has a larger number of cells per spot than osmFISH.

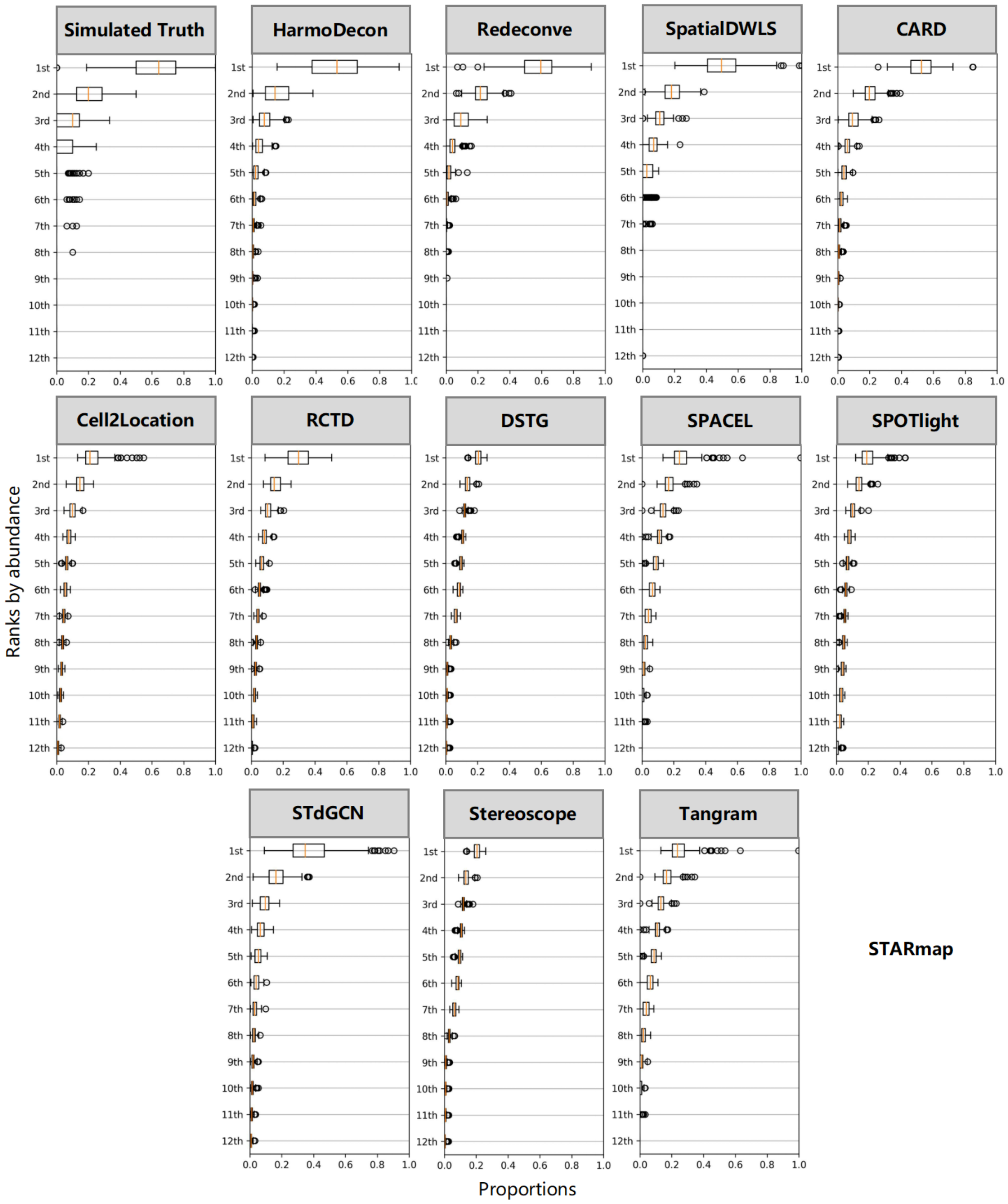
Simulated Truth
HarmoDecon
Redeconve
SpatialDWLS
CARD
Cell2Location
RCTD
DSTG
SPACEL
SPOTlight
STdGCN
Stereoscope
Tangram
STARmap
Ranks by abundance
1st
2nd
3rd
4th
5th
6th
7th
8th
9th
10th
11th
12th
Proportions
0.0
0.2
0.4
0.6
0.8
1.0

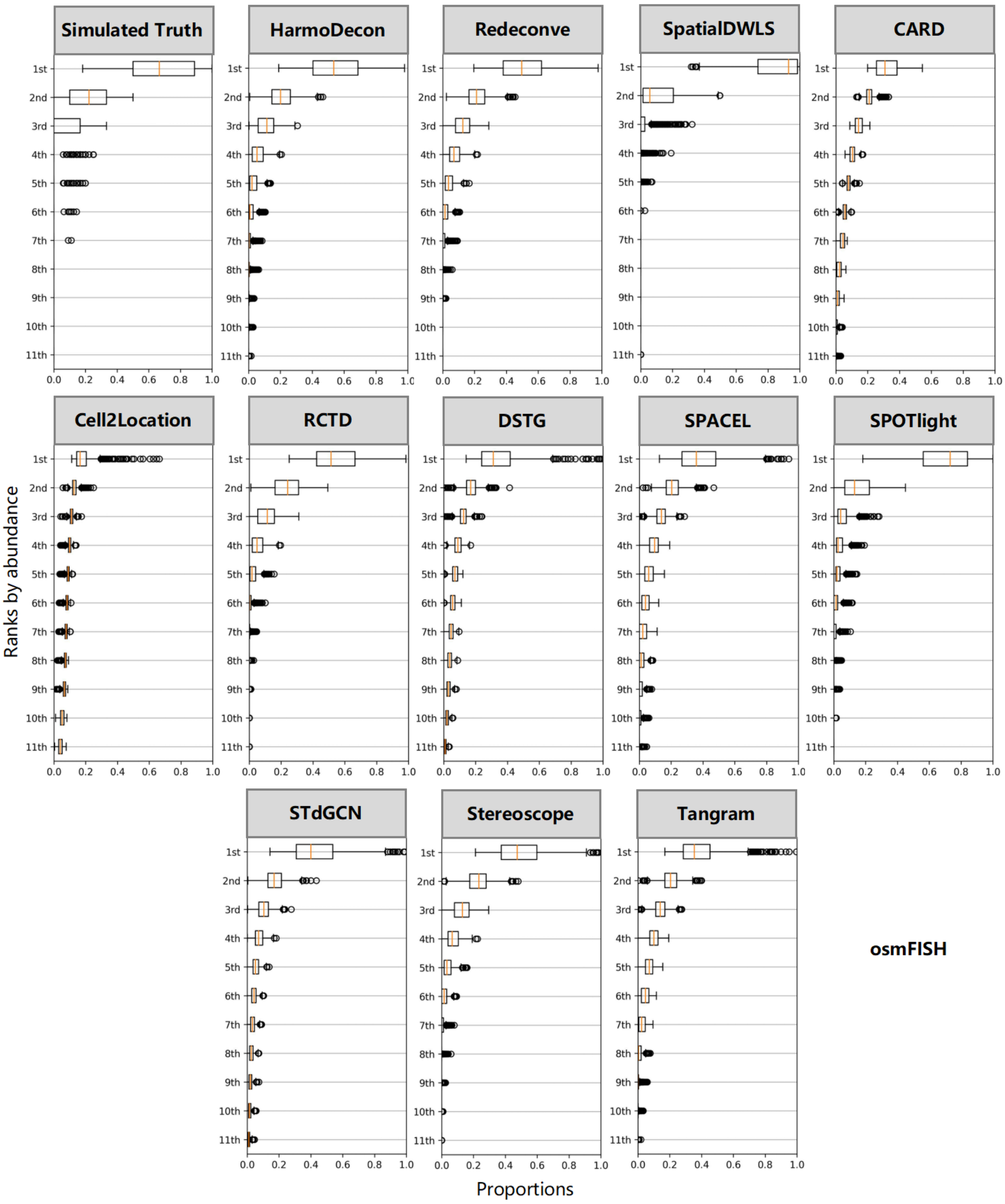


**Supplementary Figure 2. The box plot of cell-type proportions ordered by abundance per spot (n = 189) in the simulated STARmap[1] and osmFISH data[3].** We obtained the simulated 'multi-cell spots' STARmap and osmFISH data with cell-type annotations from previous studies[2,4]. We sorted the proportions in each simulated multi-cell spot of the observed truth and results inferred by 12 deconvolution methods. From the most significant proportion values to the last, we displayed their distributions by boxplots.

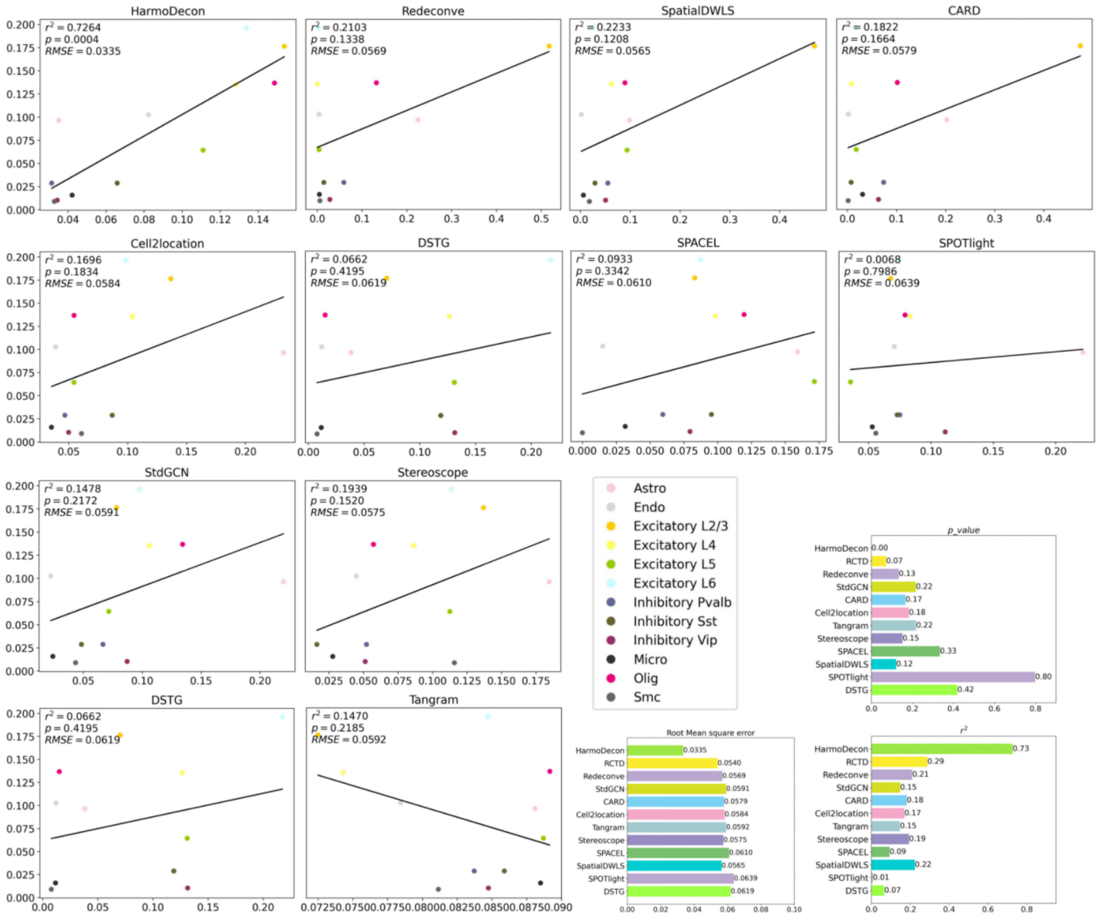

HarmoDecon
$r^2 = 0.7264$
$p = 0.0004$
$RMSE = 0.0335$
Redeconve
$r^2 = 0.2103$
$p = 0.1338$
$RMSE = 0.0569$
SpatialDWLS
$r^2 = 0.2233$
$p = 0.1208$
$RMSE = 0.0565$
CARD
$r^2 = 0.1822$
$p = 0.1664$
$RMSE = 0.0579$
Cell2location
$r^2 = 0.1696$
$p = 0.1834$
$RMSE = 0.0584$
DSTG
$r^2 = 0.0662$
$p = 0.4195$
$RMSE = 0.0619$
SPACEL
$r^2 = 0.0933$
$p = 0.3342$
$RMSE = 0.0610$
SPOTlight
$r^2 = 0.0068$
$p = 0.7986$
$RMSE = 0.0639$
StdGCN
$r^2 = 0.1478$
$p = 0.2172$
$RMSE = 0.0591$
Stereoscope
$r^2 = 0.1939$
$p = 0.1520$
$RMSE = 0.0575$
DSTG
$r^2 = 0.0662$
$p = 0.4195$
$RMSE = 0.0619$
Tangram
$r^2 = 0.1470$
$p = 0.2185$
$RMSE = 0.0592$
Astro
Endo
Excitatory L2/3
Excitatory L4
Excitatory L5
Excitatory L6
Inhibitory Pvalb
Inhibitory Sst
Inhibitory Vip
Micro
Olig
Smc
p_value
HarmoDecon 0.00
RCTD 0.07
Redeconve 0.13
StdGCN 0.22
CARD 0.17
Cell2location 0.18
Tangram 0.22
Stereoscope 0.15
SPACEL 0.33
SpatialDWLS 0.12
SPOTlight 0.80
DSTG 0.42
Root Mean square error
HarmoDecon 0.0335
RCTD 0.0540
Redeconve 0.0569
StdGCN 0.0591
CARD 0.0579
Cell2location 0.0584
Tangram 0.0592
Stereoscope 0.0575
SPACEL 0.0610
SpatialDWLS 0.0565
SPOTlight 0.0639
DSTG 0.0619
$r^2$
HarmoDecon 0.73
RCTD 0.29
Redeconve 0.21
StdGCN 0.15
CARD 0.18
Cell2location 0.17
Tangram 0.15
Stereoscope 0.19
SPACEL 0.09
SpatialDWLS 0.22
SPOTlight 0.01
DSTG 0.07


#### STARmap

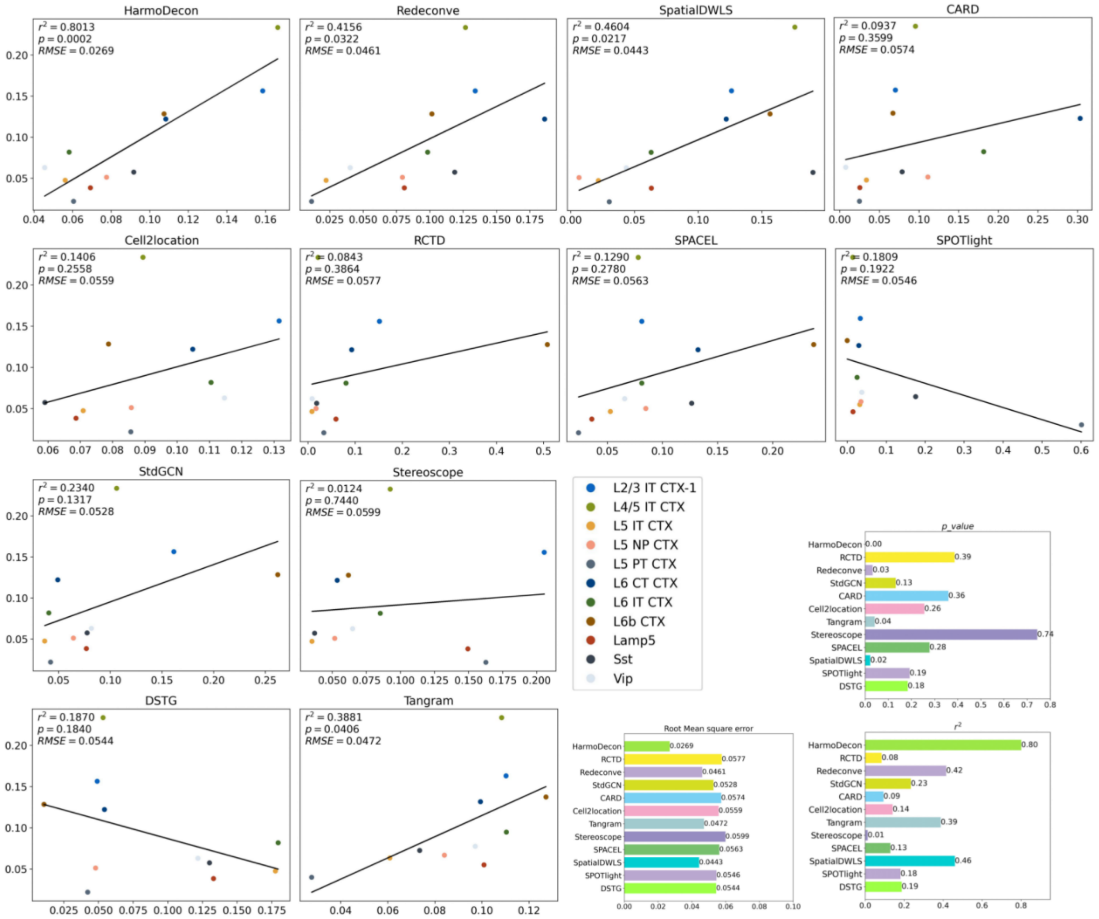


##### osmFISH

**Supplementary Figure 3. HarmoDecon successfully captures the linear correlation when recovering the sample-level cell-type profiles.** Apart from demonstrating superior performance in inferring spot-level cell-type proportions, HarmoDecon consistently delivered promising results when summing the proportions of spots to reconstruct sample-level cell-type profiles, showing a high correlation. When examining the p-value for a hypothesis test utilizing the Wald Test with a t-distribution of the test statistic, only HarmoDecon consistently showed significance($p < 0.05$) across the two single-cell spatial transcriptomics datasets. Our method effectively captured the linear relationship in sample cell-type fractions.

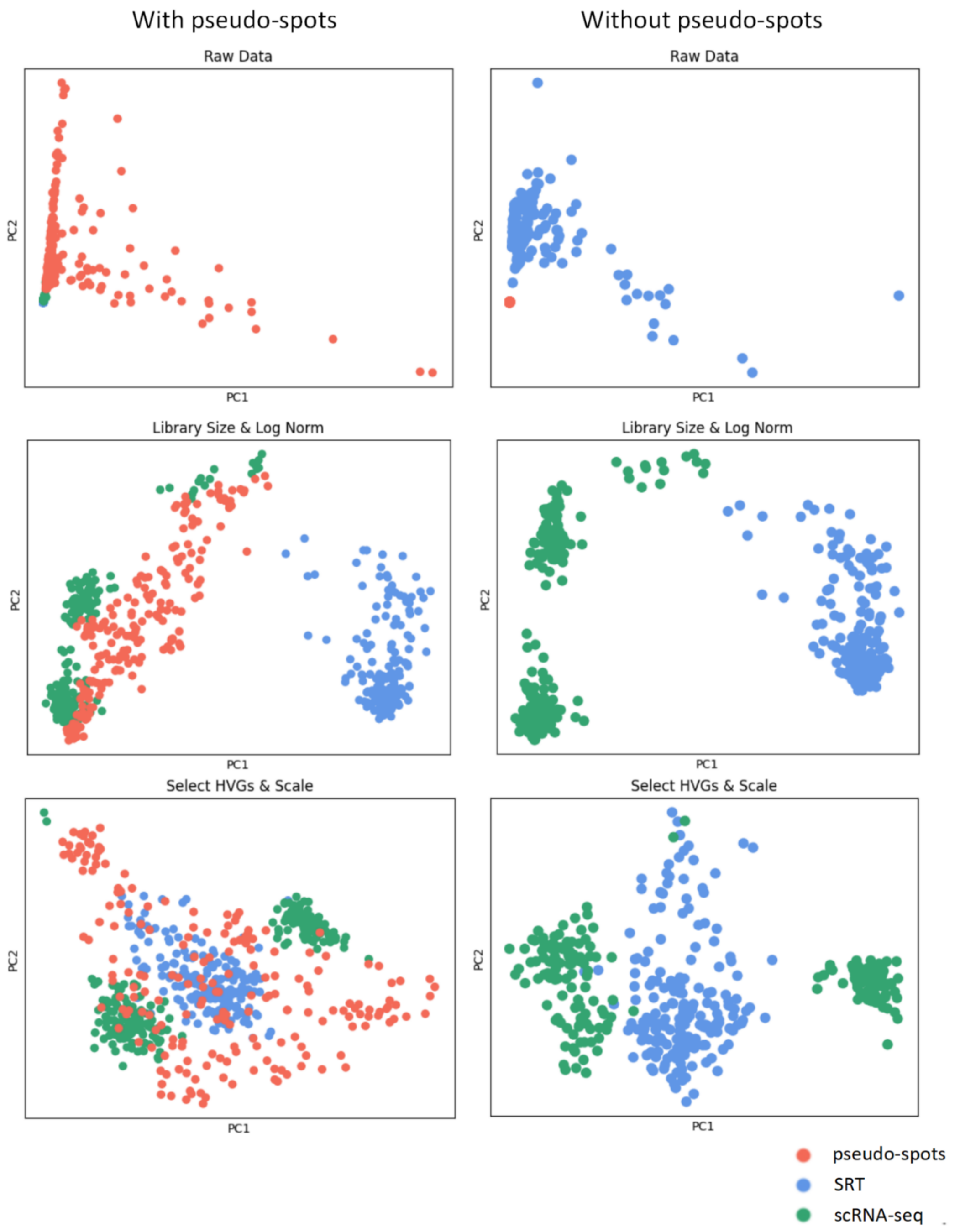


**Supplementary Figure 4. Pre-processing can alleviate the platform effects that occur in the STARmap dataset.** By looking at the scatter plots after PCA, we observed that the raw gene expression of scRNA-seq (green points) and SRT simulated multi-cell spots (blue points) has a strong deviation. After library size and logarithmic normalization, the raw gene expression of scRNA-seq and SRT points still do not have an overlapping area. As mentioned in the paper, our method generated pseudo-spots by aggregating the expression values of scRNA-seq. The synthesized pseudo-spots have an overlapping area with the target SRT data used for deconvolution.

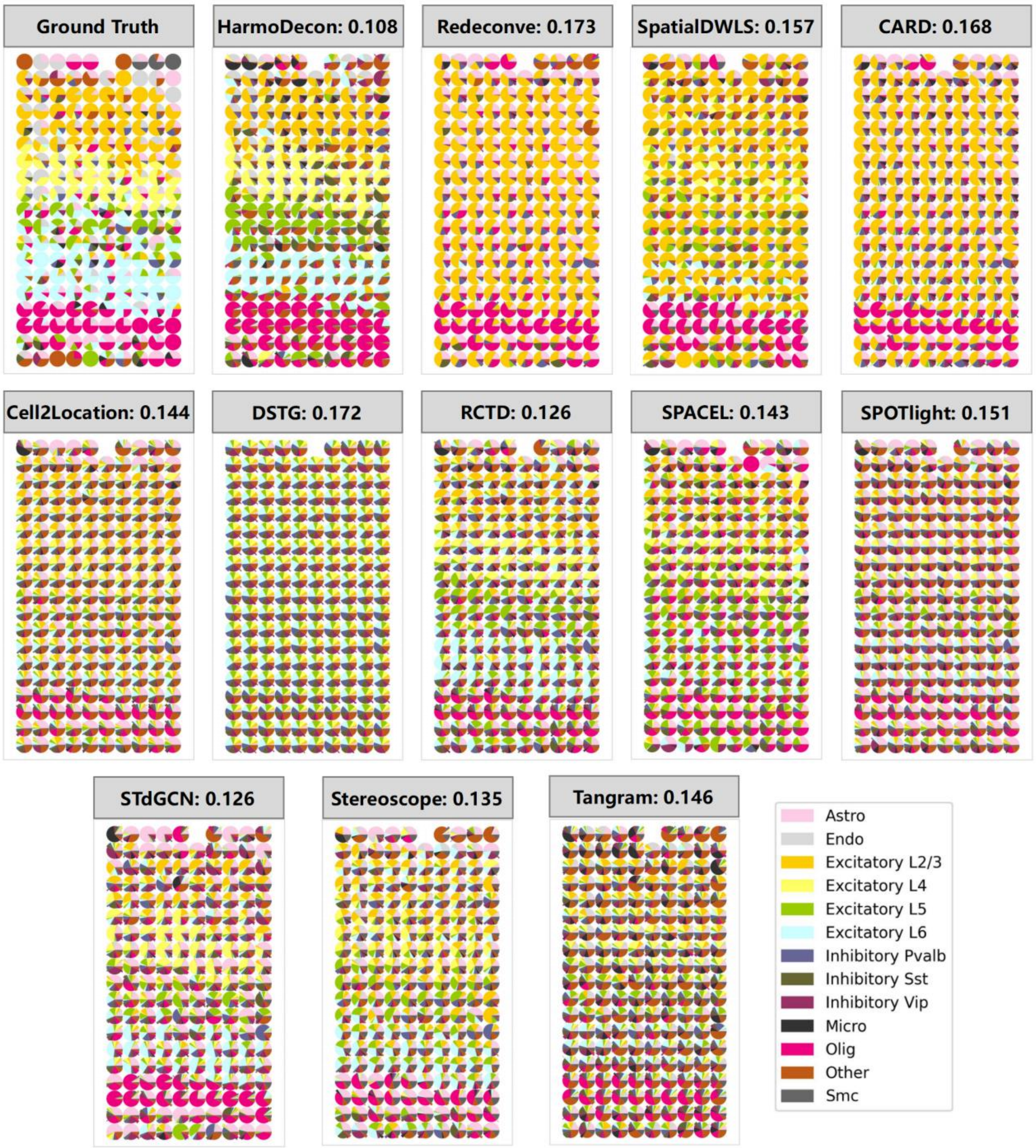


**Supplementary Figure 5. Pie chart of the deconvolution results of STARmap mouse visual cortex data.** While the article only presents a subset of the results from selected well-performing methods, we have included the complete results here as a supplement. Our observations indicated that HarmoDecon consistently outperforms other methods in terms of the evaluated Root Mean Square Error (RMSE).

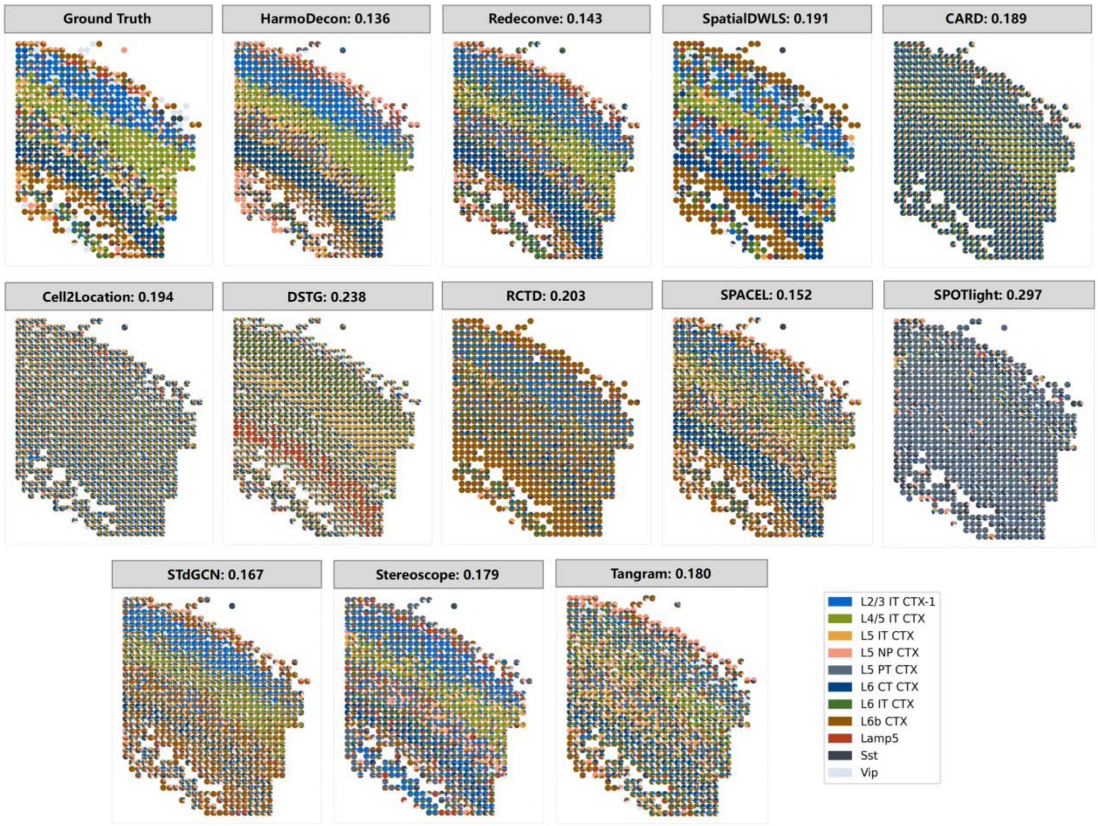


**Supplementary Figure 6. Pie chart of the deconvolution results of osmFISH mouse somatosensory data.** While the article only presents a subset of the results from selected well-performing methods, we have included the complete results here as a supplement. Our observations indicated that HarmoDecon consistently outperforms other methods in terms of the evaluated Root Mean Square Error (RMSE).

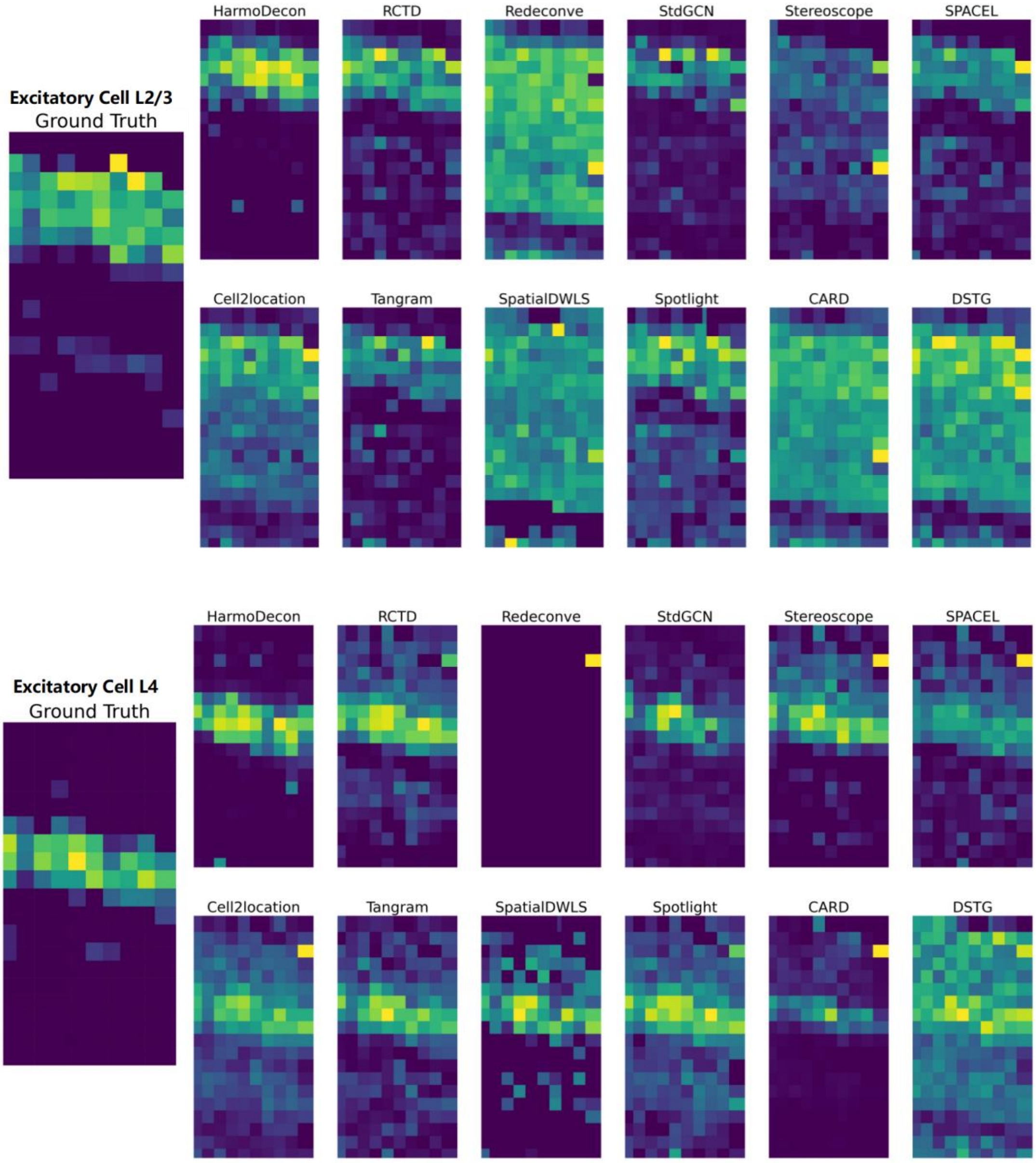
Excitatory Cell L2/3
Ground Truth
HarmoDecon
RCTD
Redeconve
StdGCN
Stereoscope
SPACEL
Cell2location
Tangram
SpatialDWLS
Spotlight
CARD
DSTG
Excitatory Cell L4
Ground Truth
HarmoDecon
RCTD
Redeconve
StdGCN
Stereoscope
SPACEL
Cell2location
Tangram
SpatialDWLS
Spotlight
CARD
DSTG

145

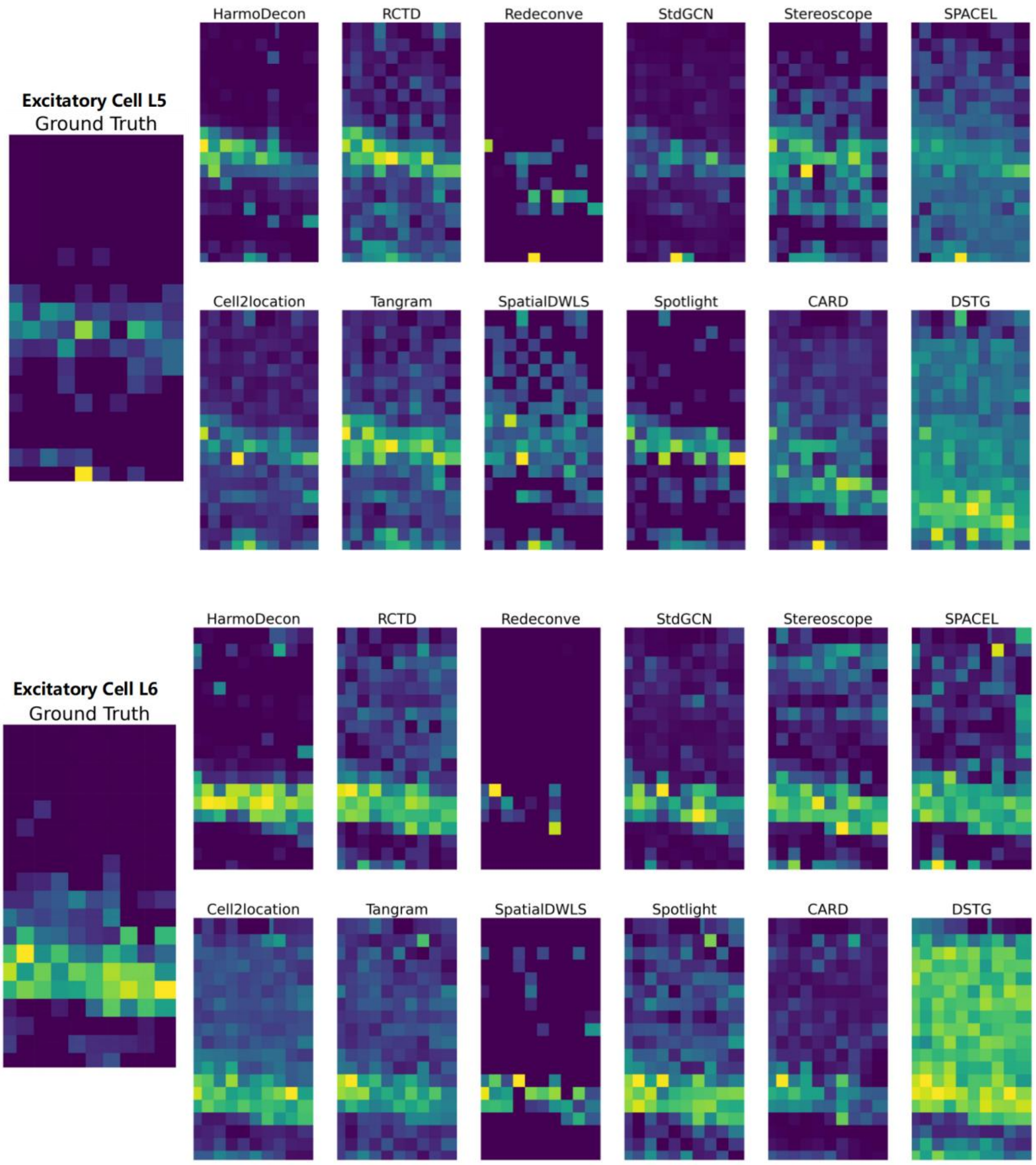


**Supplementary Figure 7. Visualization of main cell types in STARmap dataset.** We have chosen several layer-specific excitatory cells for demonstration purposes. As illustrated in the figure below, only HarmoDecon, RCTD, and StdGCN consistently exhibited a clear boundary for each anatomical layer. Among these methods, HarmoDecon stood out by having fewer false positive spots compared to the others across the four types of excitatory cells.

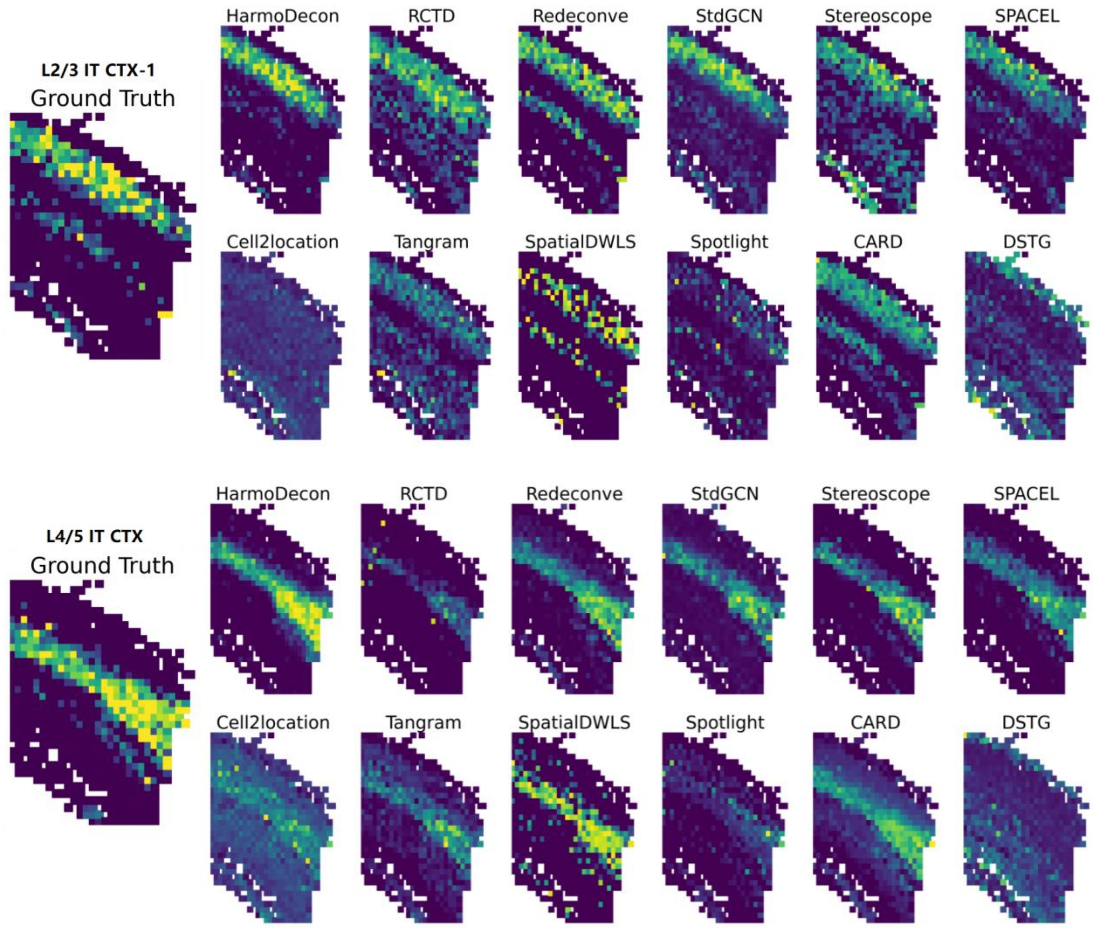
L2/3 IT CTX-1
Ground Truth
HarmoDecon
RCTD
Redeconve
StdGCN
Stereoscope
SPACEL
Cell2location
Tangram
SpatialDWLS
Spotlight
CARD
DSTG
L4/5 IT CTX
Ground Truth
HarmoDecon
RCTD
Redeconve
StdGCN
Stereoscope
SPACEL
Cell2location
Tangram
SpatialDWLS
Spotlight
CARD
DSTG

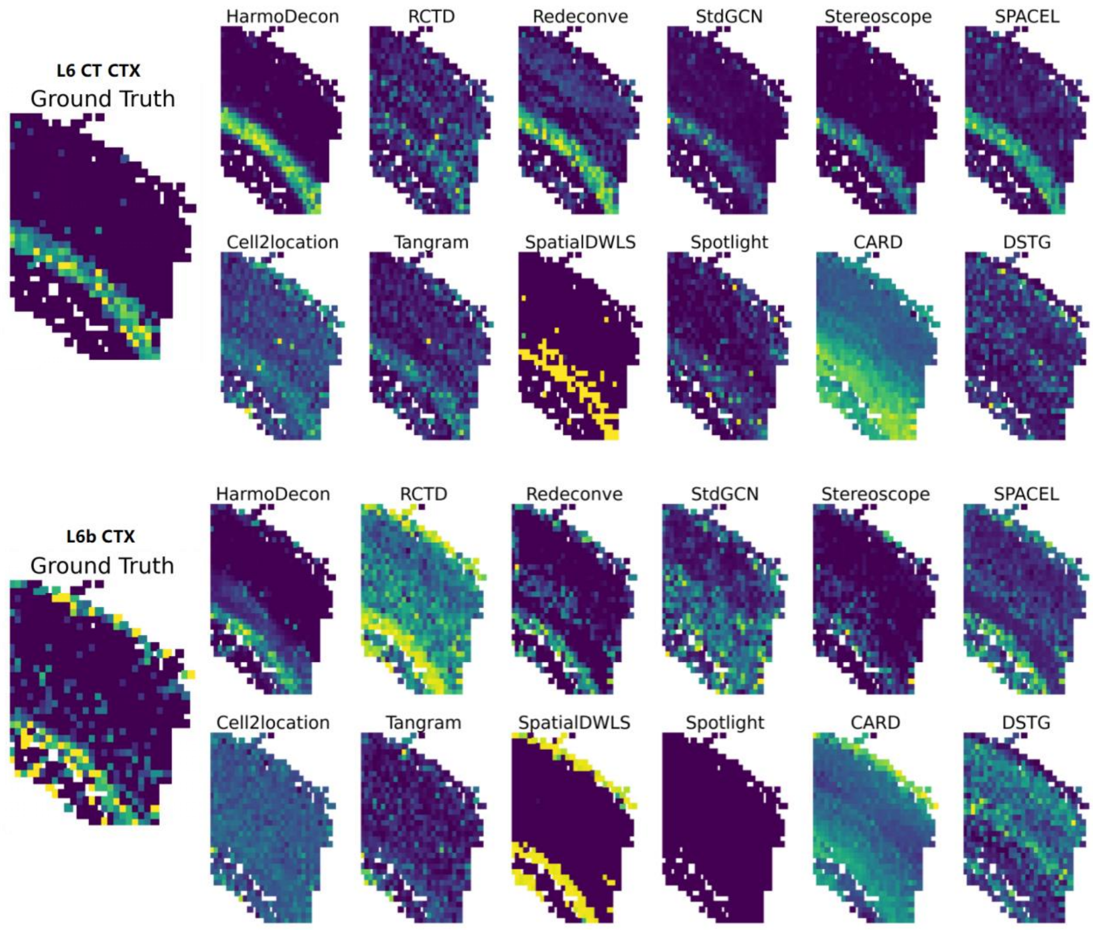


**Supplementary Figure 8. Visualization of main cell types in osmFISH dataset.** We have chosen several layer-specific excitatory cells for demonstration purposes. As illustrated in the figure below, only HarmoDecon and Redeconve consistently exhibited a clear boundary for each anatomical layer. Among these methods, HarmoDecon stood out by having fewer false positive spots compared to the others across the four types of neuron cells.

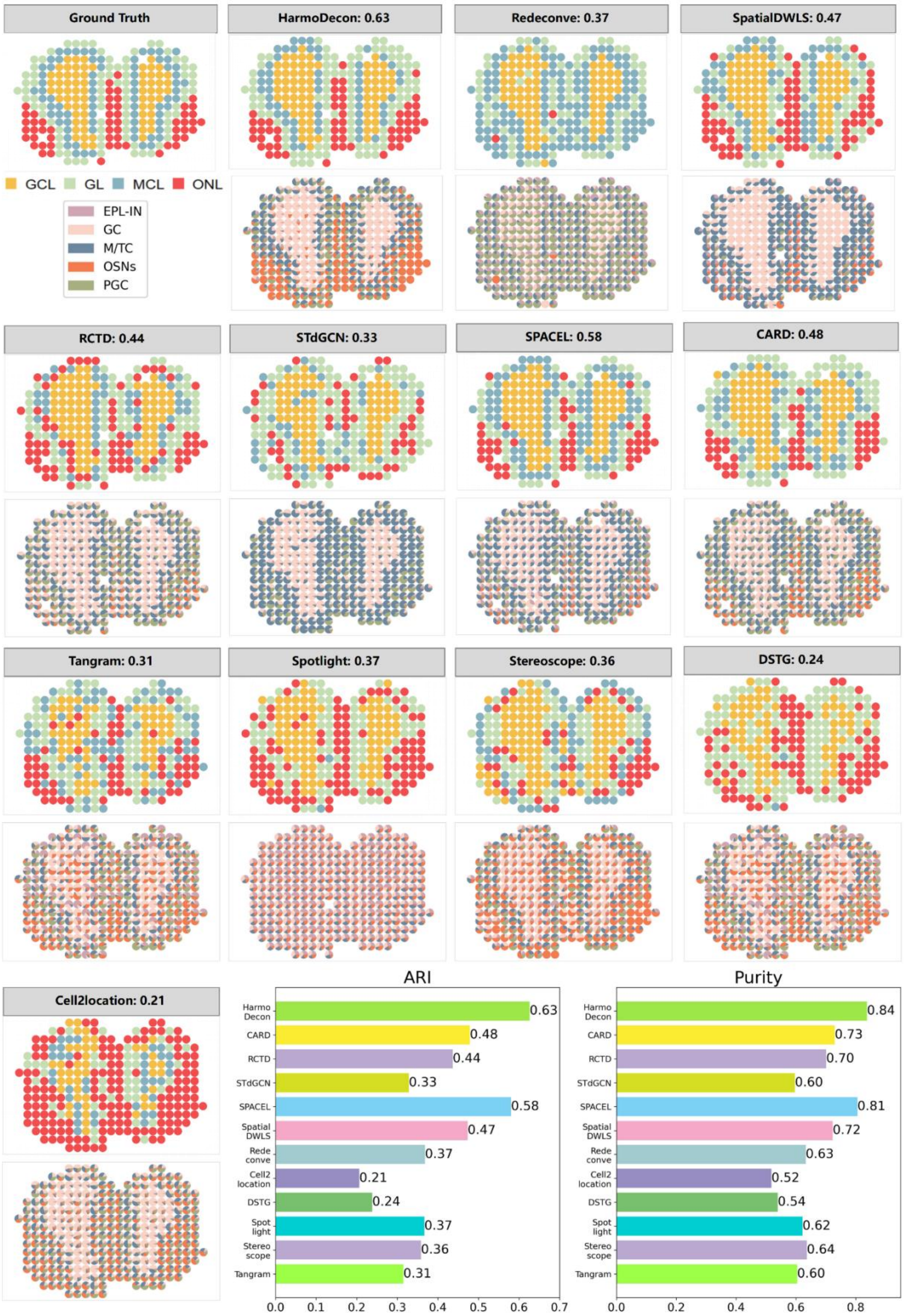


**Supplementary Figure 9. Pie charts and domain clustering of the deconvolution results of mouse olfactory bulb (MOB)[6] data by legacy ST.** These figures demonstrated that HarmoDecon consistently surpasses other methods in the evaluated Adjusted Rand Index (ARI) and purity with domain clustering by cell types.

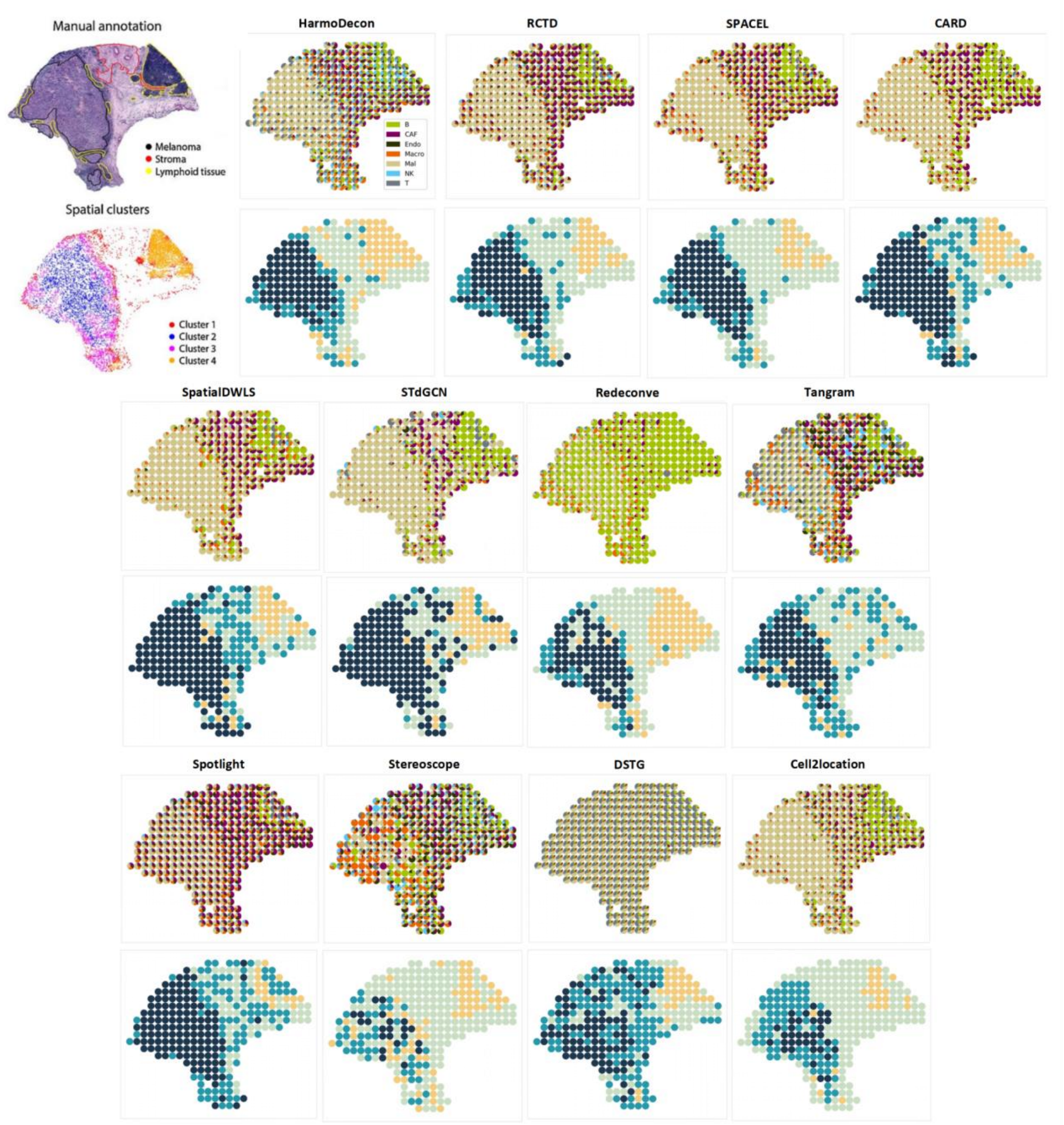


**Supplementary Figure 10. Pie chart and domain clustering of the deconvolution results of human melanoma[7] data by legacy ST.** Figures show that only HarmoDecon, RCTD, and SPACEL can well distinguish the core melanoma region (dark blue-green) and border melanoma region(light blue-green).

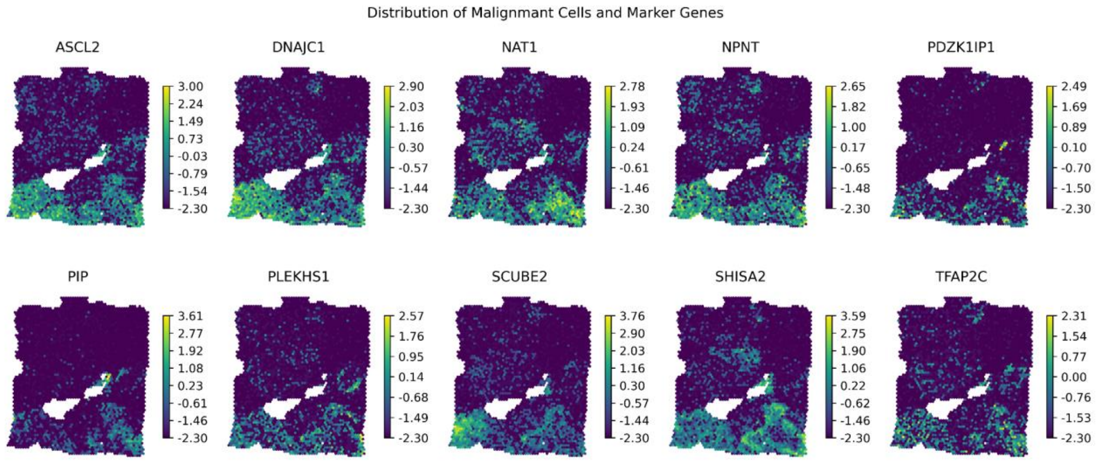


**Supplementary Figure 11. Heatmaps of selected 10 marker genes of the BRCA_1 dataset.**

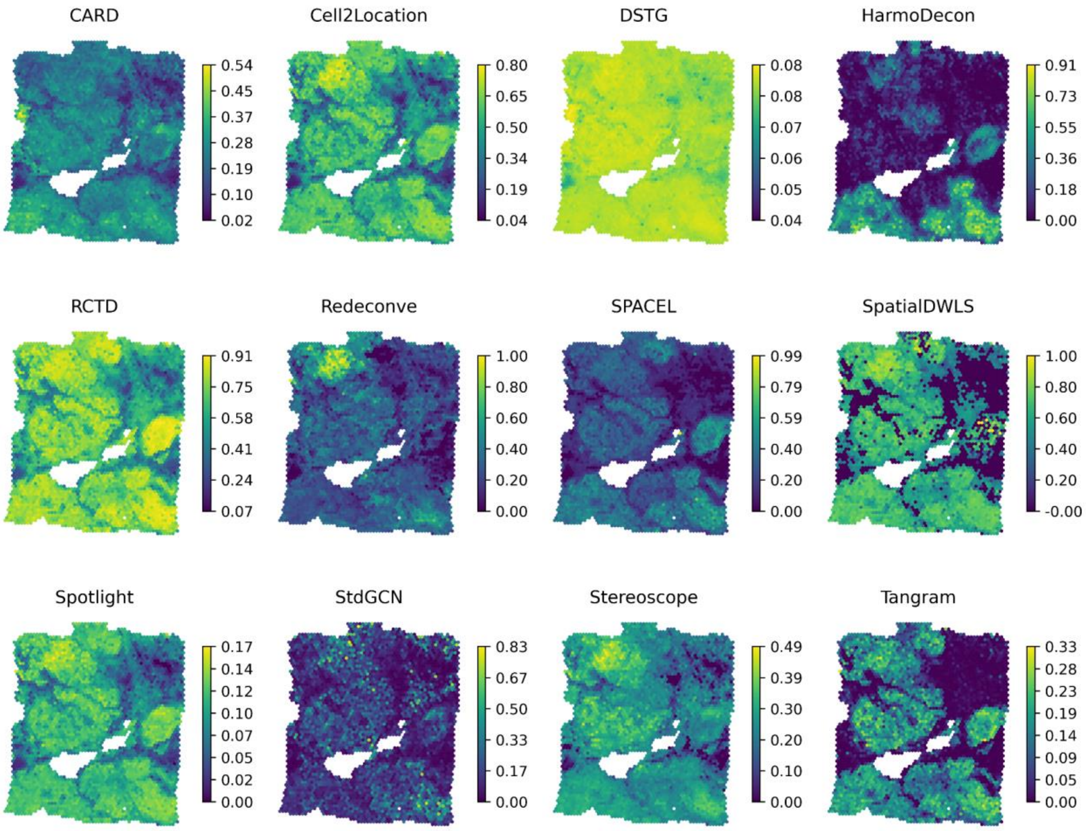


**The proportion of cancer epithelial cells inferred by 12 methods**

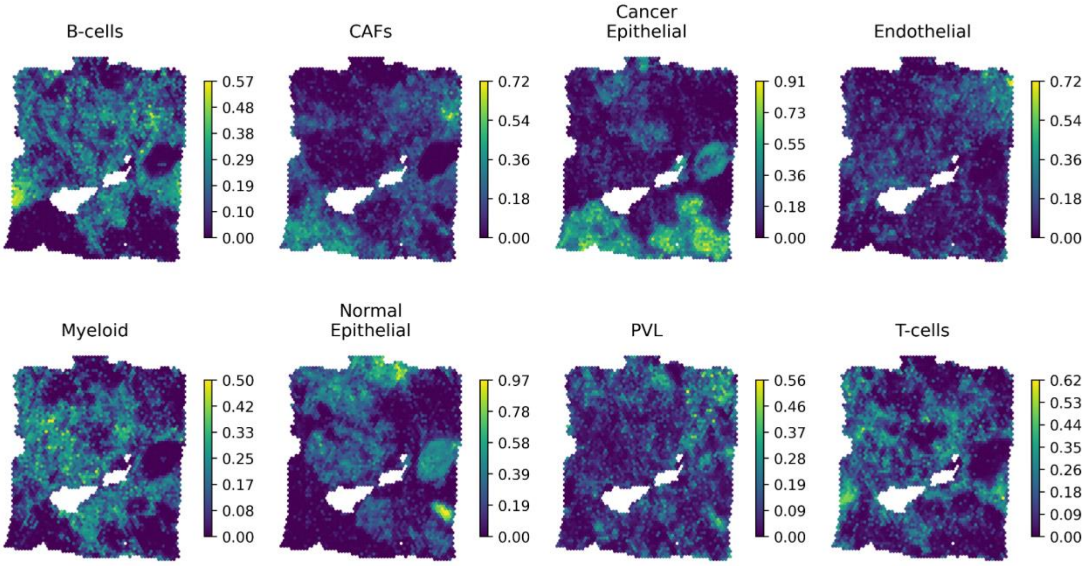


**The Proportion of each cell type inferred by HarmoDecon**

**Supplementary Figure 12. Heatmaps of the proportion of cancer epithelial cells in the BRCA_1 dataset inferred by all 12 methods.**

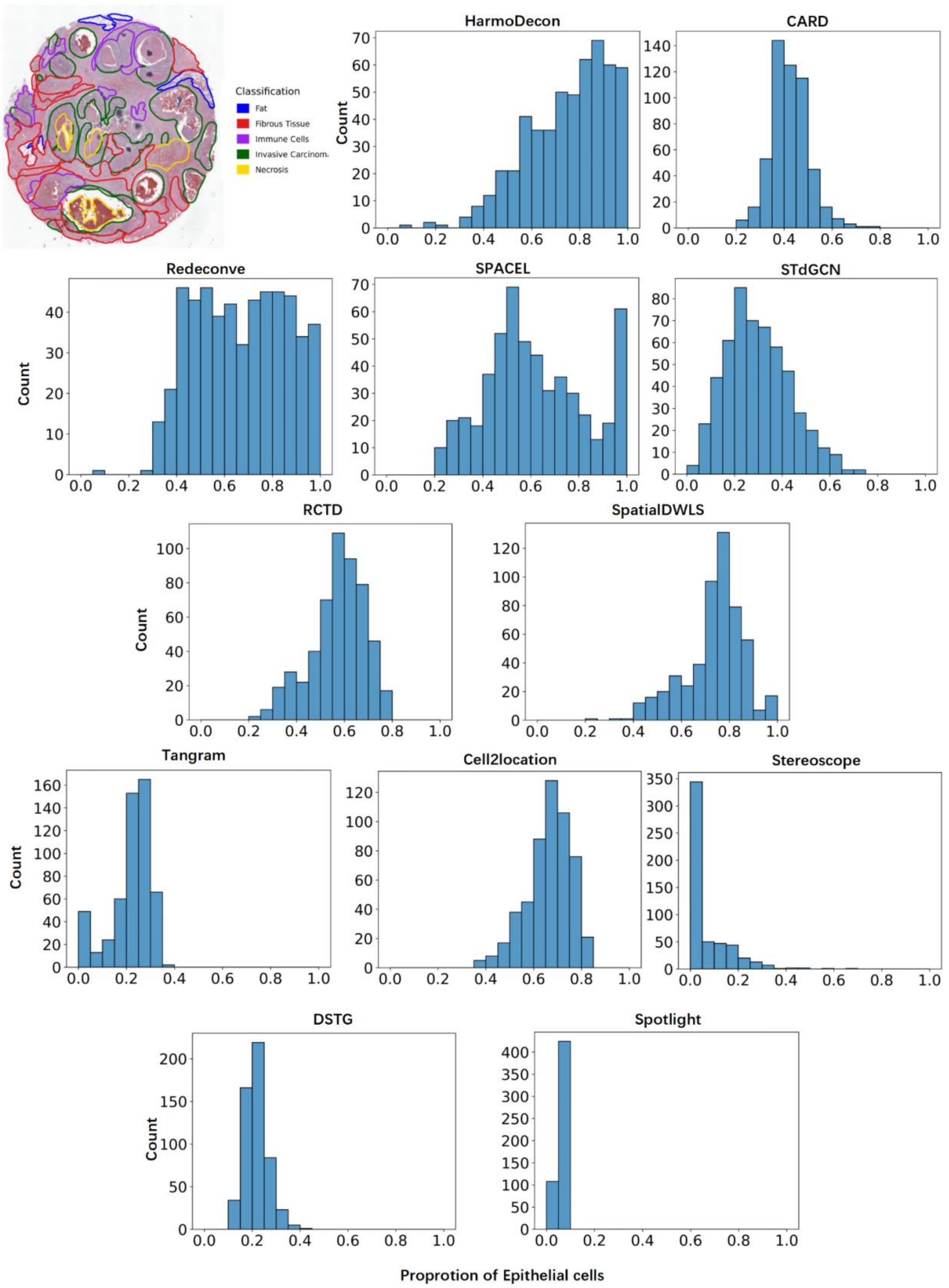


**Supplementary Figure 13. HarmoDecon captures the enrichment of epithelial cells in the invasive carcinoma region** Breast cancer lesions consist of malignant epithelial cells. In the breast cancer (BRCA_2) dataset generated by 10X Visium[5], we calculated the proportion of epithelial cells within spots located in the invasive carcinoma region (circled by green lines, 532 spots in total). Our observations revealed that among the 7 state-of-the-art methods assessed, HarmoDecon's results notably highlighted the enrichment of epithelial cells. While some modern models like Redeconve and SPACEL also captured this dominance, there were still spots within the carcinoma region that contained a lower concentration of epithelial cells.

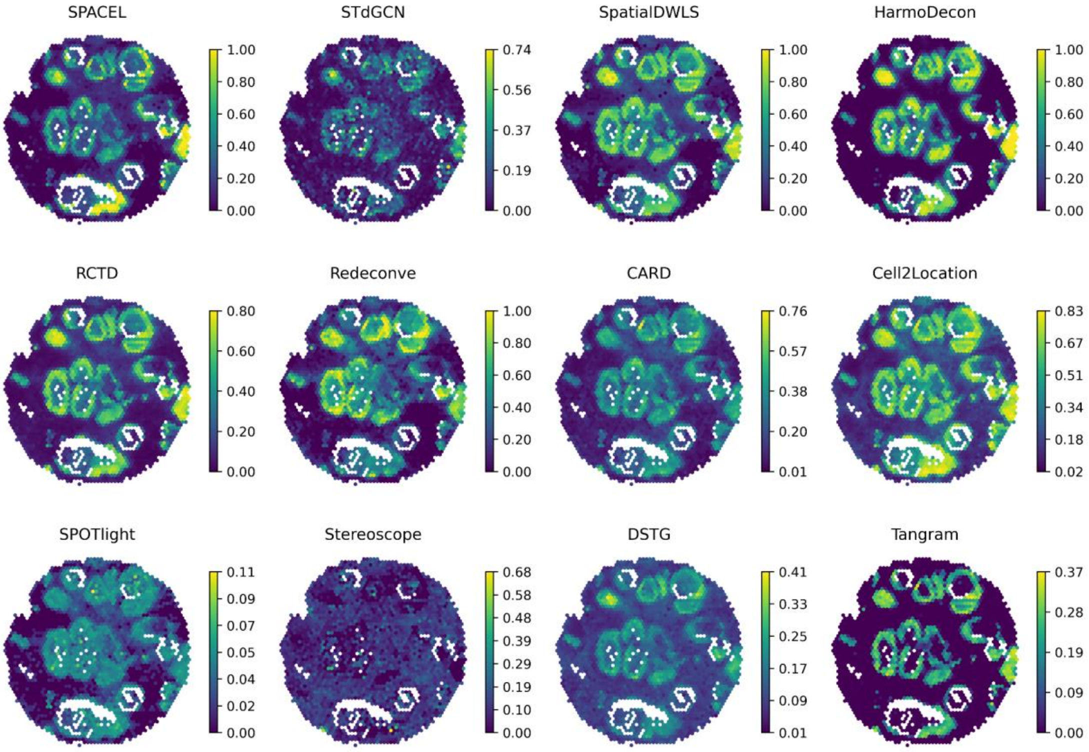


**The proportion of cancer epithelial cells inferred by 12 methods**

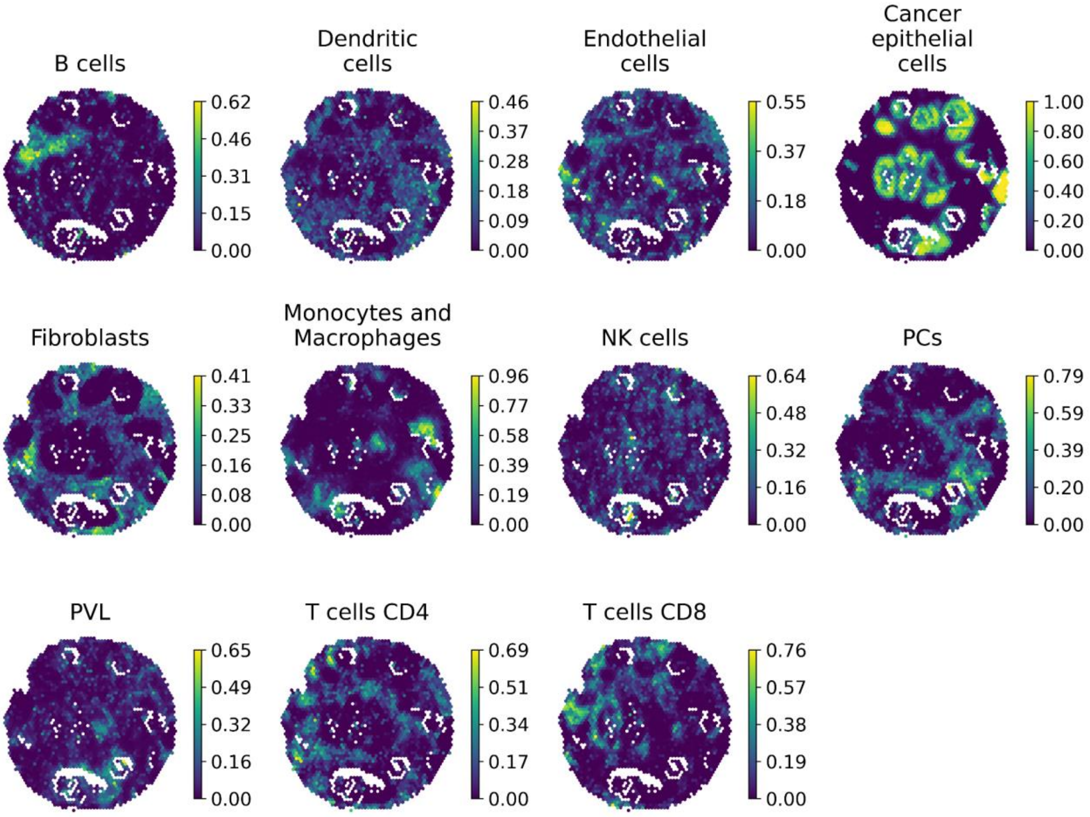


**The Proportion of each cell type inferred by HarmoDecon**

**Supplementary Figure 14. Heatmaps of the proportion of cancer epithelial cells in the BRCA_2 dataset inferred by all 12 methods.**

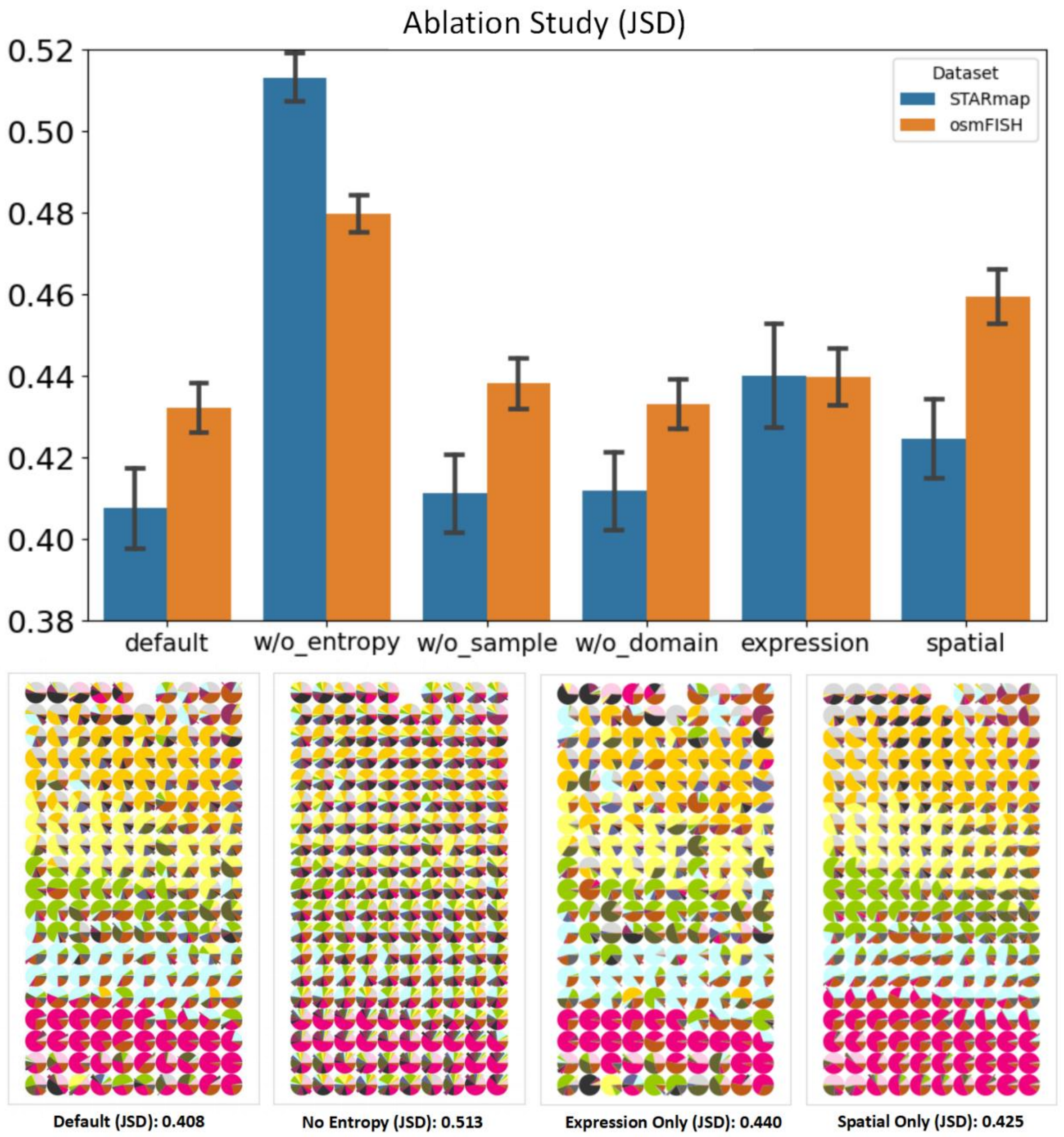


**Supplementary Figure 15. Ablation study on loss items and the graph combination strategy of HarmoDecon.** We estimated the deconvolution performances with different loss weights and the single graph on two benchmarking datasets (STARmap and osmFISH) and calculated their JS divergence compared to the ground truth. The default setting with all the loss items utilizing both the expression graph and spatial graph reaches the lowest JSD, indicating that the strategies we deployed are efficient enough to improve the deconvolution results.

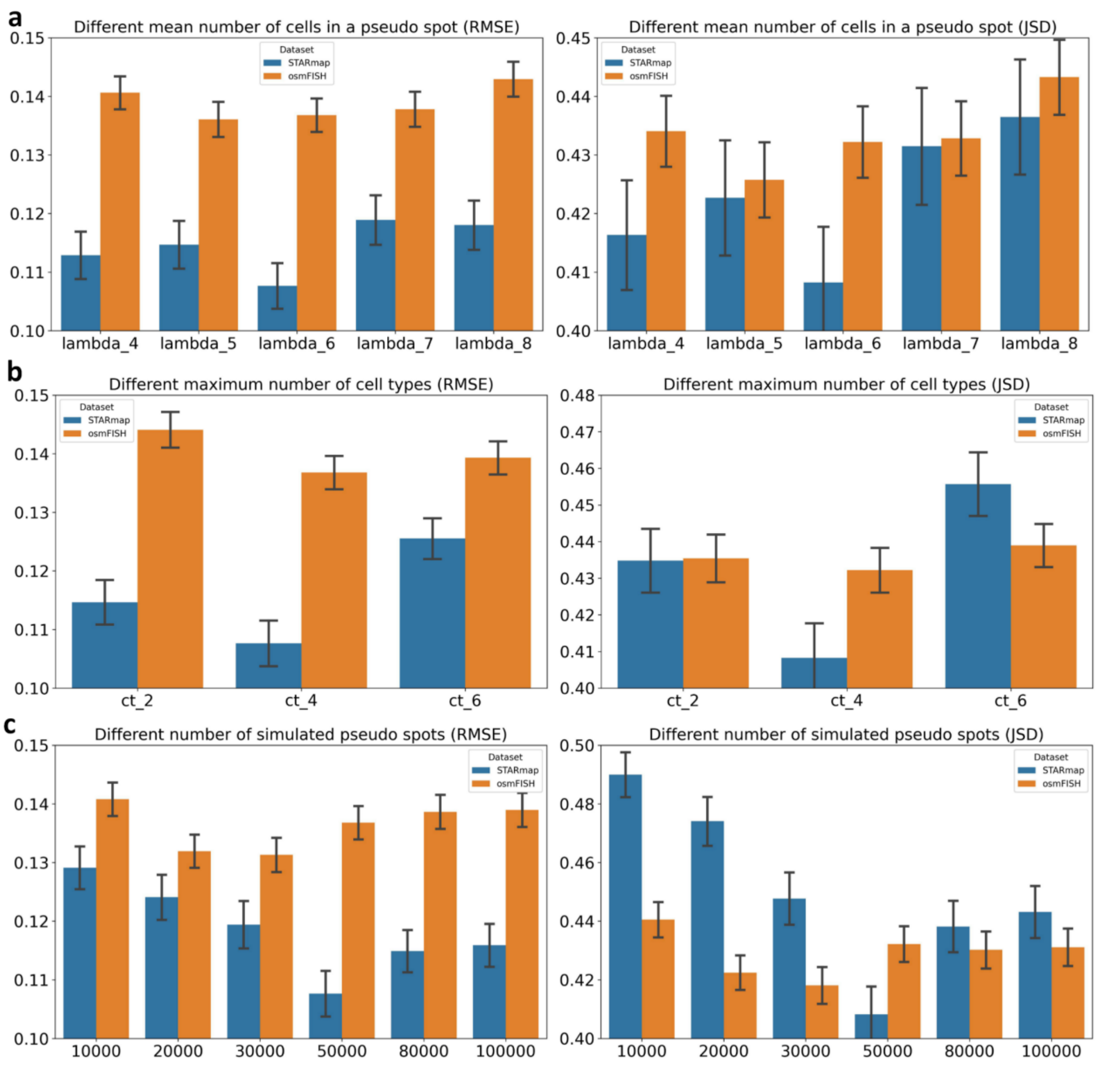


**Supplementary Figure 16. Different settings during the pseudo-spots sampling can influence the performance of HarmoDecon.** We conducted a series of ablation experiments to recommend optimal hyperparameters that are effective across various scenarios. The findings indicated that the performance was optimal when the maximum number of cell types, mean number of cells, and number of simulated spots were configured as 4, 6, and 50,000, respectively. We have selected the hyperparameters (4, 6, 50,000) as the default settings. When evaluating each parameter, the other parameters were maintained at their default values to ensure consistency and control variance.

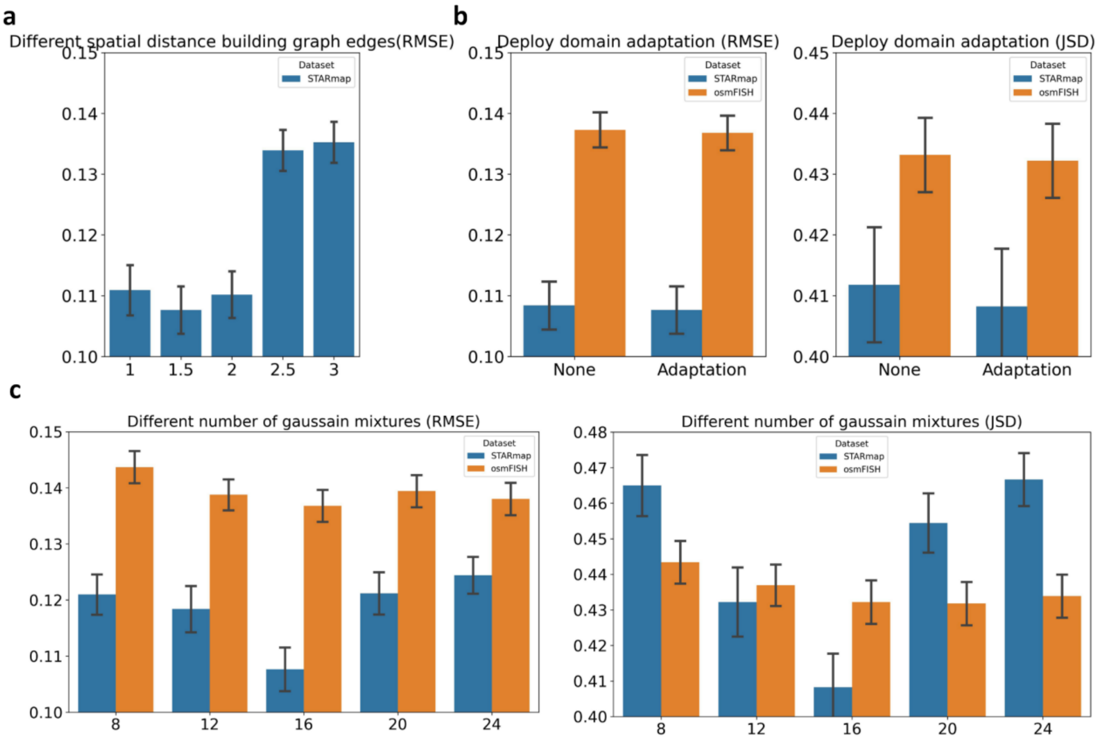


**Supplementary Figure 17. Different settings during the training period can influence the performance of HarmoDecon.** Except for the primary training loss depicted in Figure 6, we conducted additional ablation experiments to thoroughly validate the strategies implemented during the training phase. The graph on the left illustrates that varying the distance for constructing the spatial graph has a substantial impact on performance. Notably, when the distance exceeds 2, there is a significant decrease in performance. (Note: The distances calculated by coordinates [1, 1.5, 2, 2.5, 3] are approximately equivalent to K-nearest-neighbors [4, 8, 12, 20, 24]) in rectangular grids. On the right side, the figure demonstrates that integrating domain adaptation modules leads to improvements in the results, albeit not of significant magnitude.

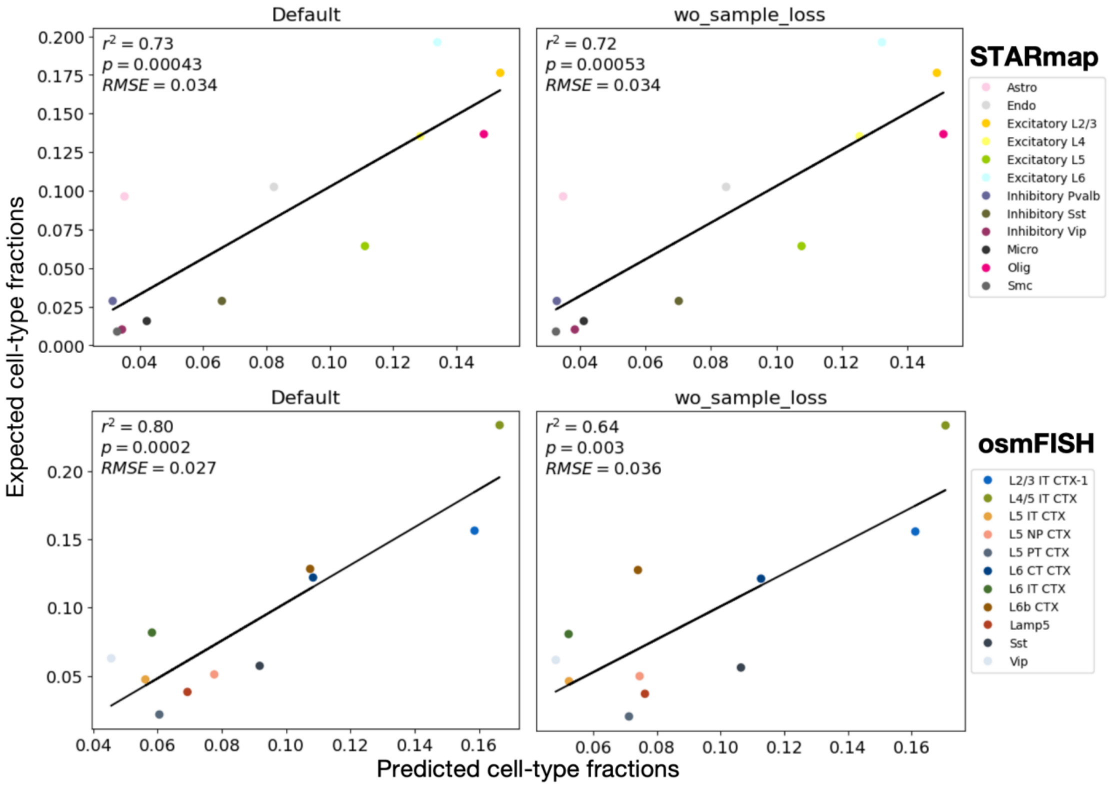


**Supplementary Figure 18. The effect of sample loss on sample-level cell-type fractions.** Left: HarmoDecon with default settings; Right: HarmoDecon without the sample loss. Upper: STARmap; Lower: osmFISH. The effect of sample loss is data-dependent. Compared to STARmap, the sample loss have more effect on the osmFISH data.

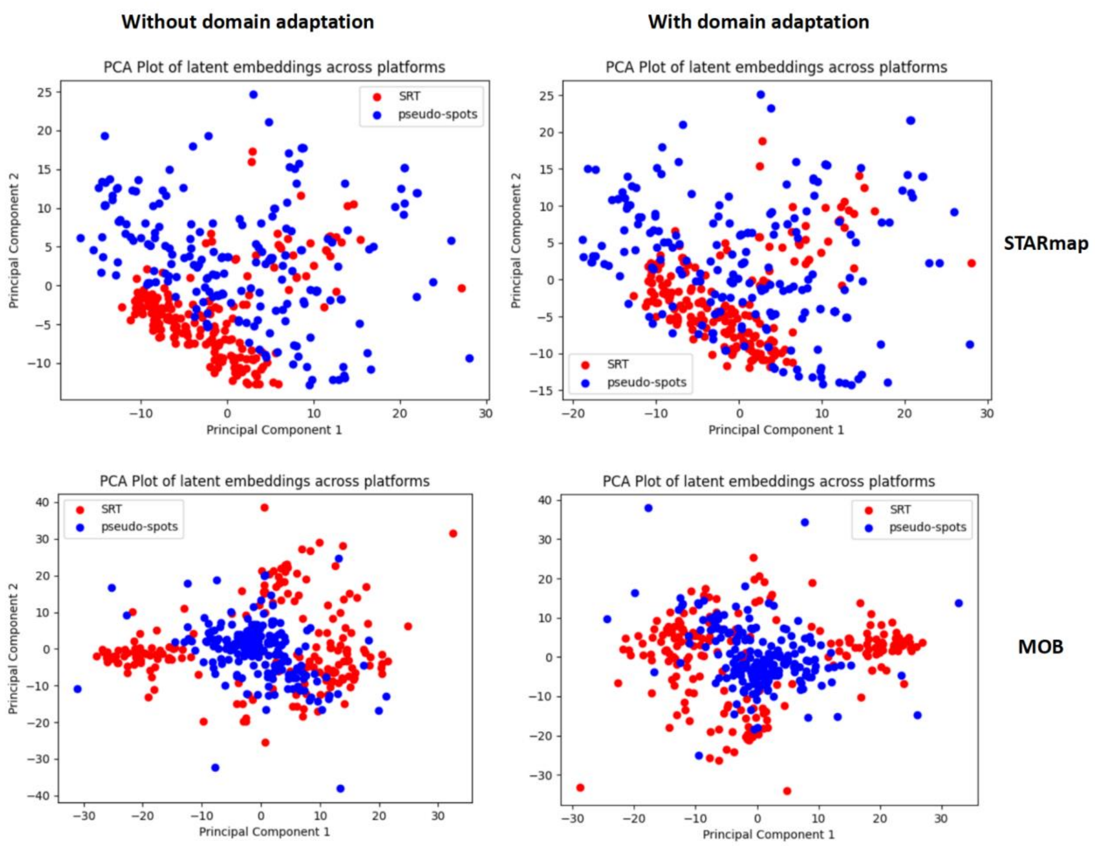


**Supplementary Figure 19. Domain adaptation module extracts common embedding vectors in the latent space.** To confirm the effectiveness of the domain adaptation module, we visualized the latent embedding vectors on STARmap and MOB datasets, projected to 2D space using PCA. We observed that after applying domain adaptation, the mixtures of 2D data points from SRT and scRNA-seq show less heterogeneity.

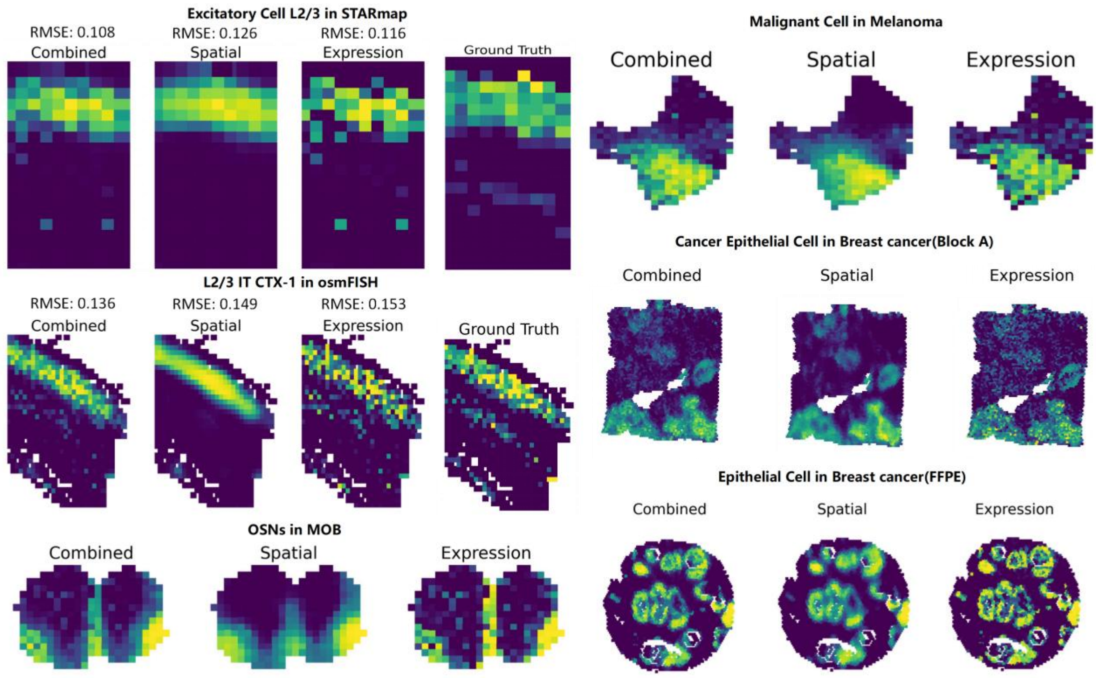


**Supplementary Figure 20. HarmoDecon combines the results inferred by the spatial neighbor graph and the expression similarity graph** In addition to gene expression profiles, HarmoDecon, being a graph-based model, accepts an extra graph as input. The edges within the graph are constructed based on either spatial neighbors or the similarity between the gene expression profiles of two spots. Across six real-world datasets, this combination consistently enhances performance. For instance, in single-cell spatial transcriptomics datasets like STARmap and osmFISH, the combined approach yields lower Root Mean Square Error (RMSE) compared to the ground truth. In bulk spatial transcriptomics datasets, the spatial component provides smoothness, while the expression component captures some domain-unseen features. For instance, in the OSNs (Olfactory Sensory Neurons) cells in the mouse olfactory bulb (MOB) dataset[6], the spatial component exhibits low false positives but may miss some border points, while the expression component contributes detailed information and detects missing values in the upper part of the olfactory nerve layer
